# Nematode Extracellular Protein Interactome Expands Connections between Signaling Pathways

**DOI:** 10.1101/2024.07.08.602367

**Authors:** Wioletta I. Nawrocka, Shouqiang Cheng, Bingjie Hao, Matthew C. Rosen, Elena Cortés, Elana E. Baltrusaitis, Zainab Aziz, István A. Kovács, Engin Özkan

**Affiliations:** Department of Biochemistry and Molecular Biology, The University of Chicago, Chicago, IL 60637, USA; Institute for Neuroscience, The University of Chicago, Chicago, IL 60637, USA; Institute for Biophysical Dynamics, The University of Chicago, Chicago, IL 60637, USA; Department of Physics and Astronomy, Northwestern University, Evanston, IL 60208, USA; Northwestern Institute on Complex Systems, Northwestern University, Evanston, IL 60208, USA; Department of Engineering Sciences and Applied Mathematics, Northwestern University, Evanston, IL 60208, USA

**Keywords:** Protein-protein interactions, Extracellular space, Cell surface receptor, Secreted protein, High-throughput interaction assay, *Caenorhabditis elegans*

## Abstract

Multicellularity was accompanied by the emergence of new classes of cell surface and secreted proteins. The nematode *C. elegans* is a favorable model to study cell surface interactomes, given its well-defined and stereotyped cell types and intercellular contacts. Here we report our *C. elegans* extracellular interactome dataset, the largest yet for an invertebrate. Most of these interactions were unknown, despite recent datasets for flies and humans, as our collection contains a larger selection of protein families. We uncover new interactions for all four major axon guidance pathways, including ectodomain interactions between three of the pathways. We demonstrate that a protein family known to maintain axon locations are secreted receptors for insulins. We reveal novel interactions of cystine-knot proteins with putative signaling receptors, which may extend the study of neurotrophins and growth-factor-mediated functions to nematodes. Finally, our dataset provides insights into human disease mechanisms and how extracellular interactions may help establish connectomes.

## INTRODUCTION

Cell surface receptors and secreted ligands build structures connecting cells, mediate communications between cells, sense and respond to extracellular cues, and function as molecular tags to identify cellular populations. These proteins include cell adhesion molecules, signaling receptors, cytokines, growth factors, and secreted cues for cellular migration among others, which are essential for most multicellular life.^1^ However, much of the extracellular proteome remains without known binding partners, which is a roadblock for revealing the various functional roles of cell surface and secreted proteins in multicellular systems. Since membrane proteins and cell surface receptors constitute the majority of targets for current therapies and are more readily accessible to new therapies,^2^ extracellular interactions are important for understanding the mode-of-action of existing therapies, as well as developing new ones.

Recent expansions of genomic sequence data and bioinformatics tools to recognize cell surface and secreted proteins have opened the study of extracellular interactions to specialized high-throughput methods.^3–5^ One of the most effective approaches has been secreted bait/prey capture strategies that use recombinantly expressed ectodomain libraries where bait and prey have been oligomerized, resulting in significant increase in effective affinity for the bait-prey pair. Such avidity-enhanced assays have been instrumental in the discovery of interactions controlling cell-cell recognition for synapse targeting and avoidance,^4,5^ neuronal repulsion,^6^ and regulatory signaling networks in plant growth and immunity,^7^ among others.

Despite these improvements, several issues with extracellular interactome studies remain: First, the existing studies have focused on an important but limited set of protein families, including the immunoglobulin (IG) superfamily^3,4,8–10^ and proteins with leucine-rich repeat (LRR) domains,^4,7,11^ while most of the known fold space in the extracellular milieu has been ignored. Second, there is a need for throughput improvements to allow for expansion of datasets towards the full interactome of cell surface and secreted proteomes. Finally, the application of the methodology to important model organisms lags. Animal genomes contain groups of conserved cell surface receptors and ligands with related functions throughout many taxa, as well as specific classes of proteins to mediate functions within few taxa or proteins with different binding partners in different taxa. The challenge of predicting whether a given set of molecular interactions are preserved across many species is a significant barrier to incorporating knowledge generated across model organisms.

The nematode *Caenorhabditis elegans* has been a pioneering model organism for studying embryogenesis, nervous system development, behavior, aging and many other aspects of biology.^12^ The *C. elegans* genome contains a smaller number of paralogs for cell surface protein families shared across bilaterians,^13^ which makes it easier to cover a larger group of receptor and ligand families relevant for human physiology. In this study, we increased the throughput of our previous avidity-based interaction screen strategy and applied it to a collection of 379 *C. elegans* ectodomains, for a putative interaction space of 72,010 possible interactions, covering 12 domain families in full and 74 domain families in part, nearly all with representation in the human genome (**Table S1**). Using a statistical method we developed for data analysis, we report 185 interactions at our intermediate confidence level, including 159 (86%) previously unknown or unpredicted by homology to known complexes. Here, we highlight and validate novel interactions between axon guidance cues and ligands from separate pathways, insulin interactions with a class of secreted IG domain proteins, and differences in interactions between mammalian and nematode orthologs. We further reveal complexes of secreted proteins of growth factor folds with their signaling receptors, and compare them to mammalian counterparts, including a novel pair with connections to human disease. Finally, we discuss interactions likely to be important in the synaptic wiring of the nervous system.

## RESULTS

### Ectodomain Library

While classical high-throughput interaction discovery technologies have proven to be highly effective for many protein classes, these strategies have remained ineffective for studying membrane proteins, cell surface receptors and secreted proteins,^14^ which make up a large fraction of animal proteomes (**Figure S1A**). Published extracellular interactome studies have focused on IG, FN (Fibronectin-type III) and LRR domains, which are some of the largest protein families in the human proteome,^3,4,7–9,11^ but have ignored the rest. To address the knowledge gap in the extracellular interactomes, we chose to study the *C. elegans* proteome (**Figure S1A**), which have smaller IG, FN, and LRR families compared to mammalian proteomes (e.g., ∼600 IG proteins in humans vs. ∼70 in C. elegans),^15,16^ allowing us to include other domain families, covering a more diverse selection of protein fold and function space with our high-throughput assay. Here, we subcloned nearly all ectodomains containing IG, FN, LRR, Cadherin (Cad), Epidermal growth factor (EGF), CUB, Laminin-N (LamN), Thrombospondin type-I (TSP1), Integrin, GAIN, Cystine-knot cytokine, and ADAM-type Cys-rich domains, as well as domain families found in known axon guidance cues and receptors (**Figures 1A-1C, Table S1**). Our ectodomain collection also includes some, but not all, ectodomains with Furin, Insulin, Kazal, Kunitz, LDLa, LY, SEA, Sushi/CCP, von Willebrand Factor type A, C and D (VWA, VWC, VWD), and WAP domains among others. Our diverse collection comprises expression constructs of 379 unique ectodomain variants, from 374 cell surface receptor and secreted protein genes with ∼86 types of domains or folds, cloned in both bait and prey expression plasmids for interaction screening, ranging from 57 to 3,220 amino acids (average: 672 amino acids), not including the tags in bait and prey expression constructs. 43% of the ectodomain library are from secreted proteins, while the rest are predicted to be cell surface receptors anchored by transmembrane helices or glycosylphosphatidylinositol (GPI) anchors (**Figure 1A**).

**Figure 1.**
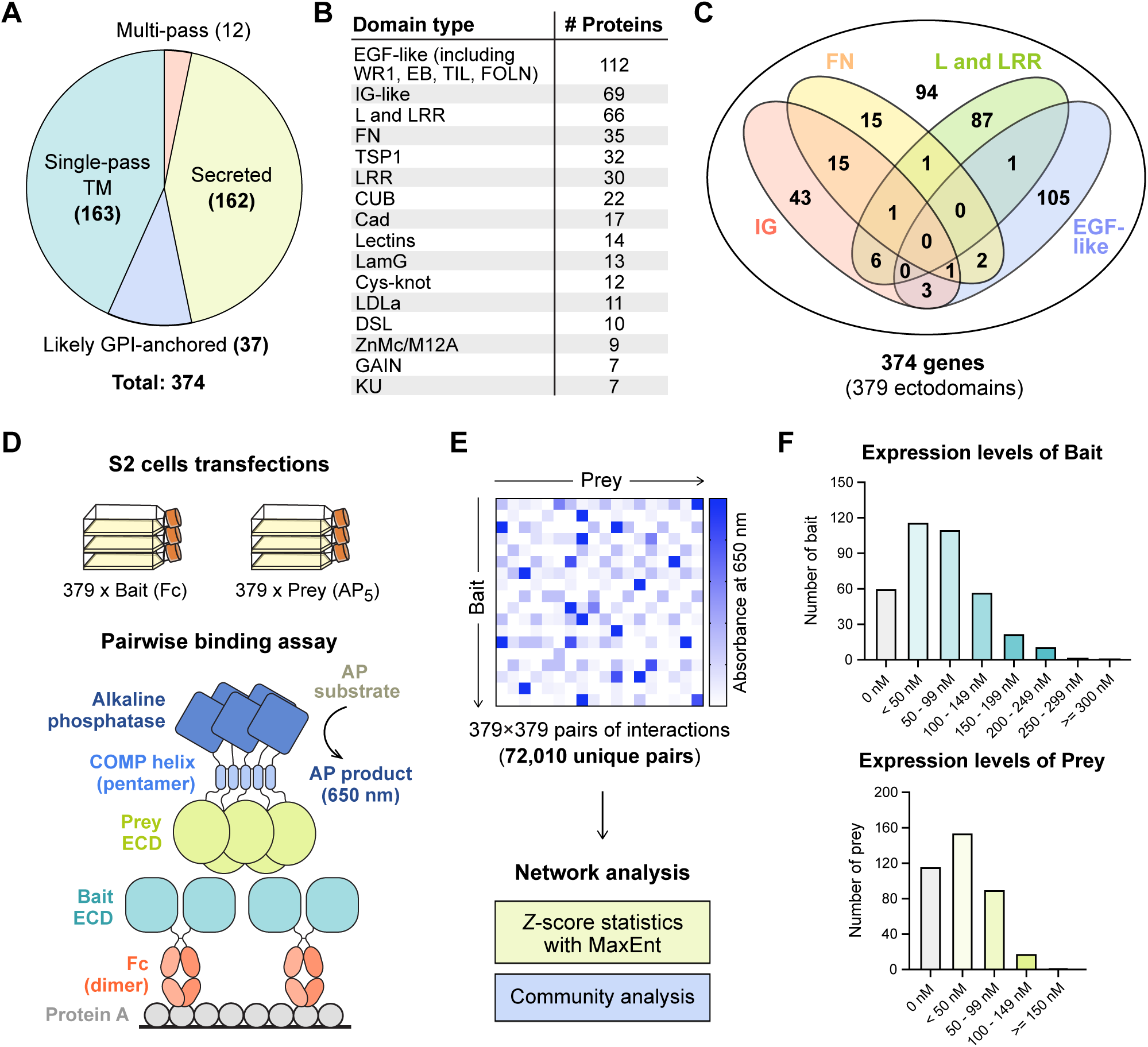
The *C. elegans* ectodomain collection and high-throughput interaction assay design. **A.** The distribution of secreted and membrane-anchored proteins in our *C. elegans* ectodomain collection. **B, C.** The distribution of protein domains in the ectodomain collection. **D, E.** The design of the ECIA pipeline and data analysis. **F.** The expression levels of bait and prey proteins in the S2 cell culture media.

### Assay Development

The Extracellular Interactome Assay (ECIA) uses Fc-tagged secreted ectodomains as bait captured on Protein A-coated plates, and Alkaline phosphatase-tagged ectodomains as prey for the detection of binding to the immobilized bait (**Figures 1D and E**). Essential to the sensitivity of the assay is the pentameric coiled coil included in the prey constructs, which increases effective affinity up to 10,000-fold through avidity,^4^ as previously implemented in similar strategies.^3,5^

The first generation of the ECIA methodology used the inducible metallothionein promoter for protein expression in the *Drosophila* S2 cell line. For this study of the *C. elegans* interactome, we continued to use the well-established S2 line, as insects are phylogenetically closer to nematodes when compared to other sources of established protein expression lines. However, we implemented several changes to simplify our protocols and improve throughput. First, we modified our expression plasmids to use the constitutively active Actin 5C promoter, which removed the induction step during expression, while improving expression levels as we observed (**Figures S1B-E**) and as previously reported for other proteins in low-throughput studies.^17,18^ We also implemented the use of 384-well plates and robotics to decrease transfection volumes, speed up the interaction assay, and decrease cost. Lastly, the advent of Gibson assembly protocols for subcloning allowed us to simplify our previous cloning strategy of Topo TA cloning followed by Gateway recombination. We cloned 63 ectodomain open-reading frames using existing cDNAs in a *C. elegans* ORF Clone Collection (GE Healthcare), 16 from our laboratories’ collections, 9 using RT-PCR from a mixed *C. elegans* mRNA library, and corrected any mutation(s) and/or intron(s) likely introduced during RT-PCR. We had the remaining 288 ORFs synthesized, choosing the longest splice variant whenever possible and practical. As we kept the Gateway recombination sequences intact in our plasmids, our ectodomain cDNA collection can be easily re-purposed for other formats and assays. Despite the improvements, 23% of our constructs did not yield detectable expression and secretion, as judged by western blotting of conditioned media (**Figure 1F**). The median concentrations in the media were estimated to be 57 nM for bait and 17 nM for prey constructs.

### Analysis of ECIA Results

ECIA reports interactions by measuring absorbance from the product of the alkaline phosphatase reaction. To detect significant interactions, we have previously calculated *z*-scores using trimmed mean and standard deviation values for each bait, and then used these normalized values to calculate *z*-scores across each prey.^4^ To improve our detection of significant hits here, we utilized a maximum entropy network ensemble-based technique^19^ to construct a statistical model that captures the absorbance background. Significant interactions are identified by comparing the observations with the modeled background (see Methods and **Figure S2** for details) using *z*-scores, where the standard deviation quantifies systematic errors. As true PPIs are likely to be detectable regardless of whether a protein is the bait or prey, we symmetrize the z-scores as 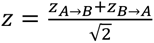 (**Table S2**). This approach yields one continuous-valued weight/score for the interaction between each pair of proteins. Using absorbance values collected at 2 hours following application of the AP substrate, an intermediate score threshold of *z_min_* = 8.4 yields a protein-protein interaction graph with 185 interactions (z>*z_min_*) (**Figures S2D, S2E** and **Table S3**). These interactions align strongly with those identified using our previously employed scoring approach (**Table S3**), but with a more rigorous standard in selecting the threshold. Lower values of *z_min_* correspond with more liberal criteria for determining which interactions to include, and thus networks with more edges.

### Community Analysis of ECIA Interaction Networks

Although protein-protein interaction networks are built from pairwise measurements, they often also reflect higher-order, functionally-relevant relationships between groups of proteins. One particularly successful approach for identifying higher-order relationships is community detection,^20^ which seeks to identify ‘insular’ subgroups of proteins – where connections between pairs of proteins within the group are stronger than those made with proteins outside the group. The interactions within these groups can reflect the sharing of latent relationships even between pairs of proteins that do not themselves directly interact. We applied a modularity optimization method to the ECIA interaction network to investigate its community structure.^21^ As expected, communities tend to group proteins with known functions (**Figure 2**). For example, proteins known to mediate axon guidance, those belonging to Robo, Slit, Ephrin and Eph, and one of the Semaphorins and Plexins, belong to the same community (community 9 in Fig. 2), while the remaining Semaphorins and Plexin are in the same community with an RPTP (PTP-4), VER-1/VEGFR, and Calsyntenin (CASY-1), proteins with known functions in regulating axon guidance (community 11 in Fig. 2).^22,23^ To capture functional groupings among a larger subset of the network (beyond the intermediate thresholds described above), we conducted the same analysis across 20-fold variation in *z_min_*, from conservative (*z_min_* = 40, *N* = 60 interactions) to liberal (*z_min_* = 2, *N* = 654 interactions). Using the communities calculated over this range of thresholds, we found nodes that repeatedly belonged to the same community. For every protein *P* for which we observed an interaction, we report (a) the full set of proteins partitioned into the same community as *P* for any tested *z_min_*, and (b) *P*’s “canonical neighbors”, the subset of proteins that belong to the same community as *P* across different choices for *z_min_*(**Table S4**).

**Figure 2.**
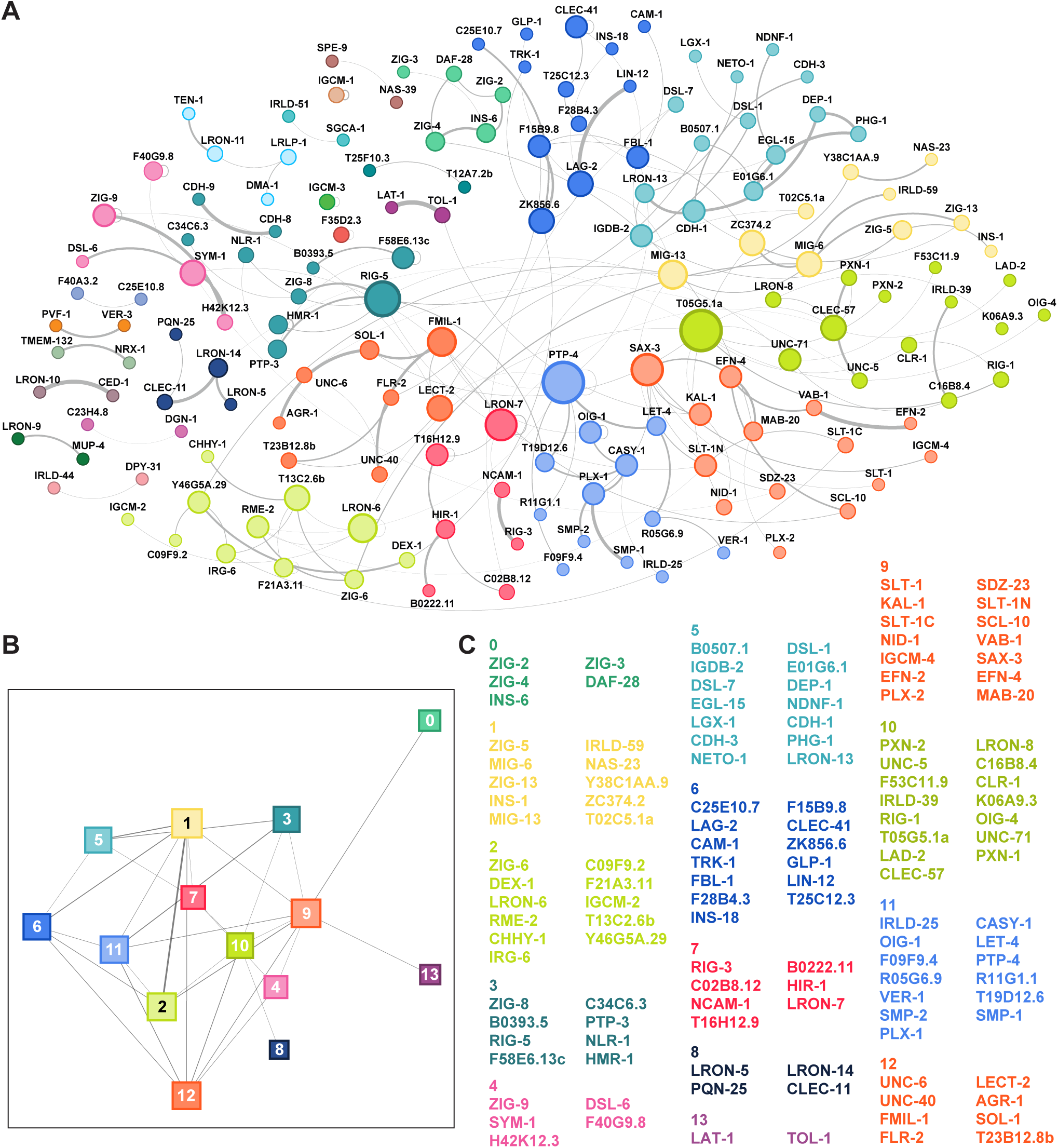
Interactions and the community structure of the extracellular interaction. **A.** Network of moderate-confidence (*z_min_*> 8.4) protein-protein interactions (*N* = 185) identified via the maximum-entropy method; proteins are colored by community grouping. **B.** Network of community-community interactions. Only *connected communities* (communities with at least one protein making at least one interaction outside the community; *N* = 14) are shown. **C.** List of proteins in each connected community.

### Higher-order axon guidance complexes

One of the highly conserved processes in nervous system development is the control of the direction of axonal growth by interactions of guidance cues with their neuronal receptors. Our study included the known cue-receptor pairs from the four classical guidance systems: Slit and Robo, Ephrins and Ephs, Semaphorins and Plexins, and Netrin (UNC-6) and its receptors, UNC-40/DCC and UNC-5.^24^ To our surprise, we observed novel interactions with high confidence that connected the three axes in the extracellular space (**Figure 3A and Table S3**). SAX-3/Robo interacts with its classical ligand SLT-1 and its N-terminal processed fragment (SLT-1N), but also with the Ephrin EFN-4 and Plexin PLX-1. This connects the Robo signaling axis to receptors of both the Ephrin and Plexin axes. In addition, EFN-4 interacts with the Semaphorin MAB-20, physically connecting Ephrin receptor with a Plexin ligand in the extracellular space. We successfully replicated these results and others involving guidance receptors and cues using ECIA (**Figures 3B and S3**), resulting in a connected network of interactions where only the Netrin axis remained unconnected in the extracellular space (**Figure 3A**). It should be noted that the DCC class of Netrin receptors were shown previously to interact with the Robo receptor intracellularly.^25^ We also identified two novel ligands for Netrin receptors: PXN-1, a peroxidasin associated with neuronal phenotypes,^26^ interacting with UNC-5; and FMIL-1, an adhesion GPCR, interacting with UNC-40/DCC.

**Figure 3.**
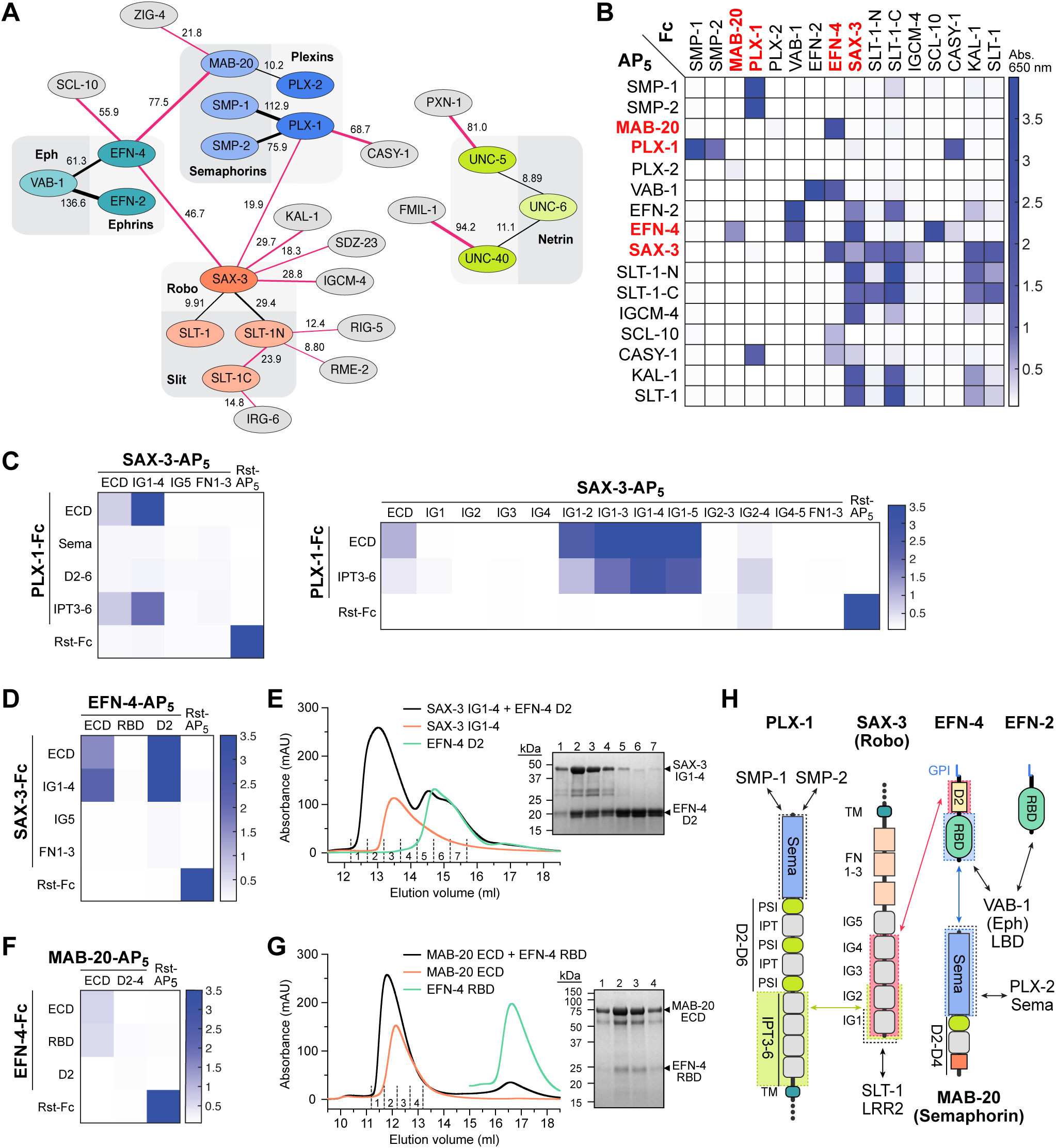
Axon guidance receptors and cues interact with each other outside the known cue-receptor axes. **A.** Schematic of interactions from the high-throughput assay, where line thickness is scaled to symmetrized MaxEnt *z*-scores (Figure 2). Interactions that were not previously known are shown with red lines. The numbers next to the lines indicate *z*-scores. **B.** Interactions observed in the high-throughput assay are reproduced (see Figure S3 for more). As expected, interactions observed with a bait-prey (Fc-AP) pair are also observed in the reciprocal orientation, resulting in diagonally symmetrical assay results. **C-G.** ECIA can be used to identify domains required for novel interactions: The first two IG domains of SAX-3 interact with the IPT domains 3 to 6 (C). SAX-3 IG domains also interact with the second domain of EFN-4, as observed by ECIA (D) and size-exclusion chromatography (SEC) (E). The interaction of MAB-20 with EFN-4 is mediated by the first domain (RBD) of EFN-4, as observed by ECIA (F) and SEC (G). SDS-polyacrylamide gels show the presence of both ectodomains in the complex fractions of SEC runs (E, G). **H.** Summary of interactions between the domains of the axon guidance receptors and cues. Black arrows refer to interactions we observed but also previously characterized in other taxa.

Given the central importance of axonal guidance pathways to neuronal wiring and their involvement in neurodevelopmental disorders, we chose to further validate these interactions with orthogonal methods. First, we performed ECIA experiments with protein domains to learn about the architectures of these new complexes. We showed that the IG domains of SAX-3/Robo bind PLX-1 IPT domains 3 to 6, and EFN-4 domain 2 (**Figures 3C and 3D**). To confirm these data, we produced SAX-3 IG1-4, EFN-4 D2 and PLX-1 IPT3-6 domain constructs using the baculoviral expression system in lepidopteran cells, and purified them to homogeneity. Purified SAX-3 IG1-4 and EFN-4 D2 can form a stable complex observed via size-exclusion chromatography (SEC) (**Figure 3E**). We also validated the interaction of the Semaphorin MAB-20 with the Ephrin EFN-4, using purified MAB-20 ectodomain with the first domain of EFN-4 with both ECIA and SEC (**Figures 3F and 3G**). Our results implicate the first domain of Ephrin/EFN-4 in interactions with its classical receptors (EPHs) and Semaphorins, and the second domain in its interactions with Robos. Interestingly, we did not observe Semaphorin or Robo interactions with the other *C. elegans* Ephrins, which suggests specialization of EFN-4 to integrate multiple cell surface signals.

### A family of insulin-receptor complexes revealed in nematodes

Insulin and related peptides are conserved peptide hormones that regulate metabolism, cell proliferation, aging and longevity across animals through a conserved set of downstream signaling molecules, starting at the cell membrane through insulin receptors (IR) and the related Insulin-like growth factor receptors (IGFR).^27–29^ In nematodes, the insulin family has expanded to a set of 40 insulin-like peptides (ILPs: INS-1 to -39 and DAF-28),^30,31^ compared to only eight in *D. melanogaster* and ten in humans. Most nematode ILPs are expressed in neurons,^32^ and they are proposed to act through the only insulin receptor, DAF-2, in *C. elegans*.^27^ Depending on the context, ILPs have been observed to be agonists or antagonists of DAF-2,^33^ which presents a mechanistic conundrum in the absence of co-ligands or co-receptors that may modulate DAF-2 signaling in response to insulin-like peptides.

To advance our understanding of insulin signaling in *C. elegans*, we included four insulins (INS-1, INS-6, INS-18 and DAF-28) in our interaction screen. We observed that two insulins, DAF-28 and INS-6, interact with ZIG-2 and ZIG-4, respectively (**Figure 2 and Table S3**). We also observed that ZIG-3 interacts with INS-6 and ZIG-5 interacts with INS-1. Multiple lines of evidence led us to classify ZIG-2 to -5 as a novel class of insulin-binding proteins distinct from other ZIGs: (1) Our sequence alignments of ZIG molecules, previously classified as neuronal surface or secreted proteins containing two-immunoglobulin domains, showed that ZIG-2 to -5 are closely related to each other and not to other ZIGs (**Figure 4A**); (2) ZIG-2 to -5 share sequence features uncommon to other IgSF proteins, especially at the linker connecting the two IG domains, including a disulfide bond (**Figure S4A**); and (3) ZIG-2 to -5 are all secreted, where most other ZIGs appear to have transmembrane helices or GPI membrane anchors (**Table S1**). Finally, ZIG-2 to -5 are co-expressed in the PVT neuron, along with ZIG-1 and -8, and have been implicated in the maintenance of axon position in the ventral nerve chord, suggesting a functional connection among these proteins.^34,35^

**Figure 4.**
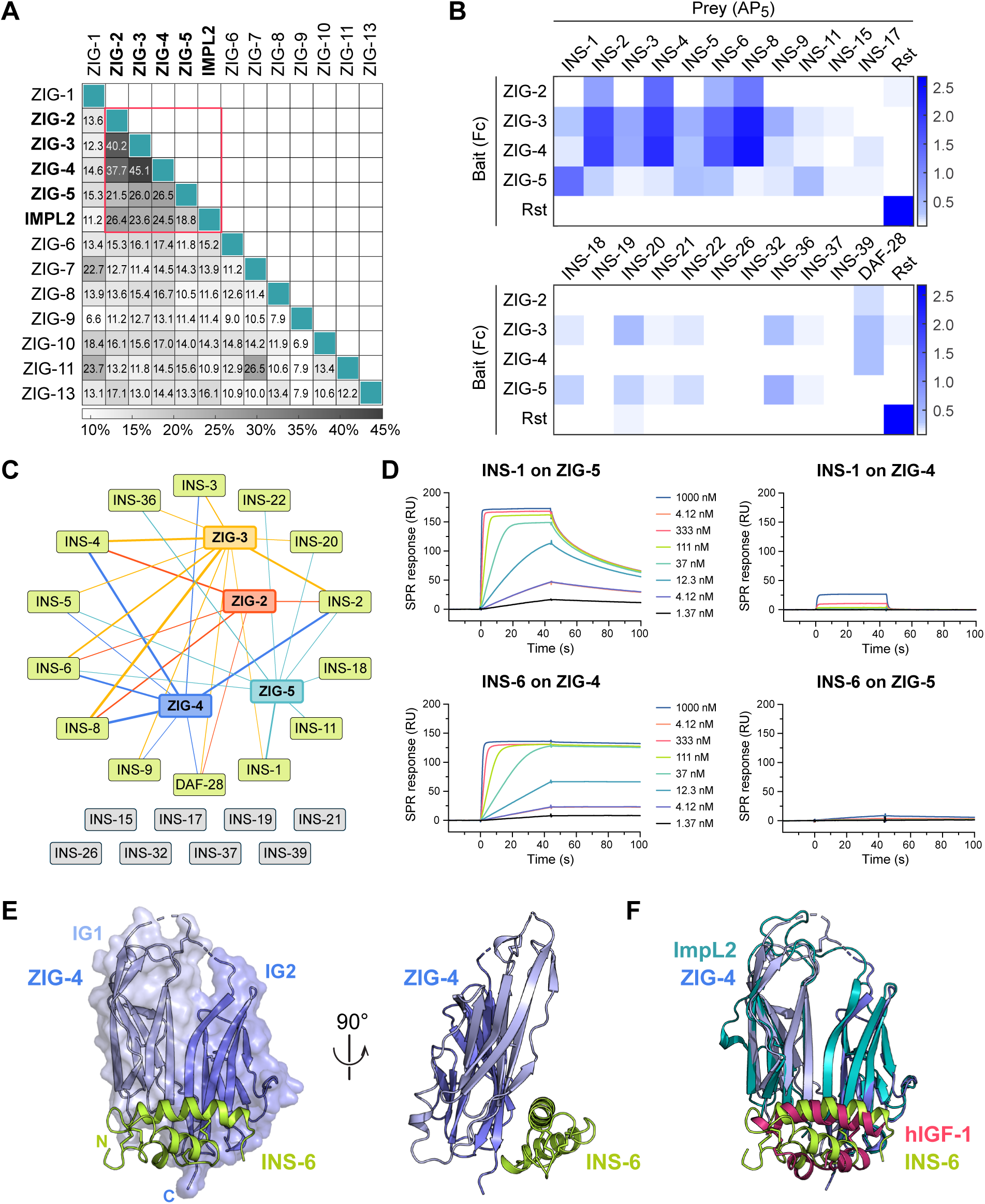
ZIG-2, -3, -4 and -5 make up a family of insulin-binding IgSF proteins. **A.** Pairwise sequence identities show that ZIG-2 to -5 are more closely related to each other and to *Drosophila* ImpL2 than the rest of the ZIG family. **B.** ECIA for 22 insulin family members against ZIG-2 to -5. For expression levels of ZIGs and insulins, see Figures S4C, S4D. **C.** ZIG-insulin interaction network inferred from the ECIA shown in B. The thickness of the connecting lines reflects the absorbance values in B for each ZIG-insulin interaction. **D.** SPR sensorgrams for INS-1 and INS-6 binding to ZIG-5 and ZIG-4 immobilized on SPR chips. Kinetic fits with estimated on- and off-rates and equilibrium constants are shown in Figure S4E. **E.** The structure of ZIG-4 (purple) bound to INS-6 (green) as observed in our tetragonal crystals. **F.** The structure of the ZIG-4–INS-6 complex strongly resembles that of *Drosophila* ImpL2 bound to human IGF-1 (PDB ID: 6FF3).

Next, we wanted to answer if other nematode insulins interact with ZIGs. To answer this question, we created ECIA expression constructs for 18 other diverse insulins via RT-PCR, covering all three classes of *C. elegans* insulins.^32^ When we repeated ECIA with these four ZIGs against the expanded set of 22 insulins, we observed binding between the four ZIGs and several other Insulins (**Figures 4B, 4C and S4B**). Since the ECIA signal depends on expression levels of bait and prey, which are different among the ZIGs and among the insulins (**Figures S4C and S4D**), we cannot compare affinities between various ZIG–Insulin interactions. However, we observed a trend where the β-class of insulins (INS-1 to -10 and DAF-28) gave higher signals of binding compared to other classes. Different binding specificities and affinities of insulins against the ZIG proteins may provide one means of differentiating their activities, even though all insulins likely act through the same receptor (DAF-2).

To validate these results, we performed SPR experiments for two pairs of insulins and ZIGs. ECIA results suggested that INS-6 binds most strongly to ZIG-4, while INS-1 binds most strongly to ZIG-5. SPR results confirmed these findings (**Figures 4D and S4E**), yielding dissociation constants (*K*_D_) for INS-1 binding to ZIG-5 and INS-6 binding to ZIG-4 were 3 nM and 56 pM, while the cross pairs had *K*_D_ > 1 µM. The very strong affinities observed suggest that insulins are unlikely to be free of ZIGs in contexts where ZIG-2 to -5 are expressed and secreted.

For more insights into the function of ZIG-insulin complexes, we crystallized the ZIG-4–INS-6 complex and determined its crystal structure in three different crystal forms (**Figure 4E**). All three structures reveal the same complex (**Figure S4F**) and show that the two immunoglobulin domains of ZIG-4 create a continuous sheet made from the *ABE* strands of IG1 and the *ACFG* strands of IG2, to which INS-6 binds (**Figure 4E**). The ZIG-4-INS-6 structure is related to recently determined structures of a *Drosophila* IgSF protein ImpL2 bound to *Drosophila* DILP5 and to human IGF-1,^36^ the atypical features of the two IG domains of ImpL2 are preserved in ZIG-4, and the overall root-mean-square displacement between the two structures is 1.4 Å over 158 (out of 202) Cα atoms (**Figure 4F**). Indeed, ZIG-2 to -5 are more closely related to ImpL2 than to other *C. elegans* ZIGs (**Figure 4A**). These similarities imply that the ancestral ecdysozoan had a ZIG-2–5/ImpL2-like protein able to interact with insulin(s). We could not identify any ZIG-4 or ImpL2 orthologs in vertebrates; however, we found ZIG-2 to -5 and ImpL2-like sequences across protostome genomes. *Drosophila* ImpL2 has been proposed as a molecule aiding the bioavailability of insulins, similar to vertebrate IGFBPs, despite sharing no ancestry, structural or sequence similarities with IGFBPs.^36^ Therefore, it is possible that ZIG-2 to -5 may be functionally related to IGFBPs in vertebrates, while not being related in sequence or structure.

As *Drosophila* and nematode insulins share the unexpected ability to act as both agonists and antagonists of insulin/IGF receptor,^31^ we analyzed how ZIG-binding to insulins could control insulin activation of the insulin/IGF receptor. We first overlaid the ZIG-4-INS-6 structure on the crystal structure of human Insulin bound to a minimal insulin receptor (IR) fragment, including the L1 domain, the Cys-rich domain, the L2 domain and the C-terminal α-helix (αCT).^37^ We saw that a ternary complex of INS-6, ZIG-4 and the insulin receptor was possible, where ZIG-4 was positioned to further interact with the IR, and not clash with it (**Figure S4G**). However, when the structure of the INS-6–ZIG-4 complex was aligned with any of the structural models of the dimeric IR bound with up to four insulins, presumably in partial or fully active states of the complex (e.g., **Figure S4H**),^38^ we observed that ZIG-4 severely clashes with one of the IR protomers forming the dimeric form of the IR complex, while aligning INS-6 with the site 1 insulin (**Figure S4I**). Similarly, the second insulin-binding site on IR overlaps with the ZIG-4-binding site, which would prevent the T-shaped IR_2_-insulin_4_ from forming (**Figure S4J**). Therefore, when ZIGs are present, insulin receptor may be able to bind insulins, but will be sequestered in an inactive form, unable to make interactions with both protomers to form an active-state IR dimer. This presents an attractive and testable mechanism for how nematode and arthropod insulins may act as antagonists of DAF-2/insulin receptor.

### Cystine-knot proteins, putative neurotrophins, growth factors and their receptors in *C. elegans*

As we curated *C. elegans* proteins that are on the cell surface or are secreted, we came across several proteins sharing common growth factor and cytokine-like folds. These include cystine-knot proteins, as well as FGF-like growth factors, and other molecules with growth factor-like sequences. We included several of these molecules in our ectodomain collection (annotated in **Table S1**) and identified binding partners for them.

First, we observed that two related cystine-knot family proteins, ZK856.6 and B0416.2, are the strongest candidates for being ligands for TRK-1 (at *z* = 22.7 and 6.3, respectively), the designated *C. elegans* ortholog of the vertebrate high-affinity neurotrophin receptors, the Trk family of receptor tyrosine kinases (RTK) (**Figure 5A**). These two secreted proteins are, therefore, putative neurotrophins, a class of growth factors necessary for neuronal growth, survival and regeneration.^39^ Vertebrate neurotrophins (NGF, BNDF, NT3 and NT4) are similarly Cys-knot proteins.^40^ Searches for homologs of the two putative neurotrophins using BLAST returned no human, mouse or fly proteins, and only each other when searched for *C. elegans* paralogs. This demonstrates that interaction screening is more effective in identifying Cys-knot ligand-receptor pairs than using sequence similarity to known pairs. To confirm our findings, we expressed and purified the TRK-1 ectodomain and one of the putative neurotrophins ZK856.6, and performed SPR experiments, which validated the interaction (**Figure 5B**). Last, structural modeling using AlphaFold-multimer showed ZK856.6 and B0416.2 dimers interacting with the FN domains of TRK-1 in a manner reminiscent of vertebrate Neurotrophin-Trk Receptor complexes (**Figure S5**), supporting our claim that TRK-1 and its interactors may signal and function similarly to neurotrophin-receptor systems in vertebrates. Interestingly, one of the putative neurotrophins, ZK856.6, is a hub also interacting with C25E10.7, F15B9.8, LAG-2 (a Jagged/Delta homolog) and DEX-1.

**Figure 5.**
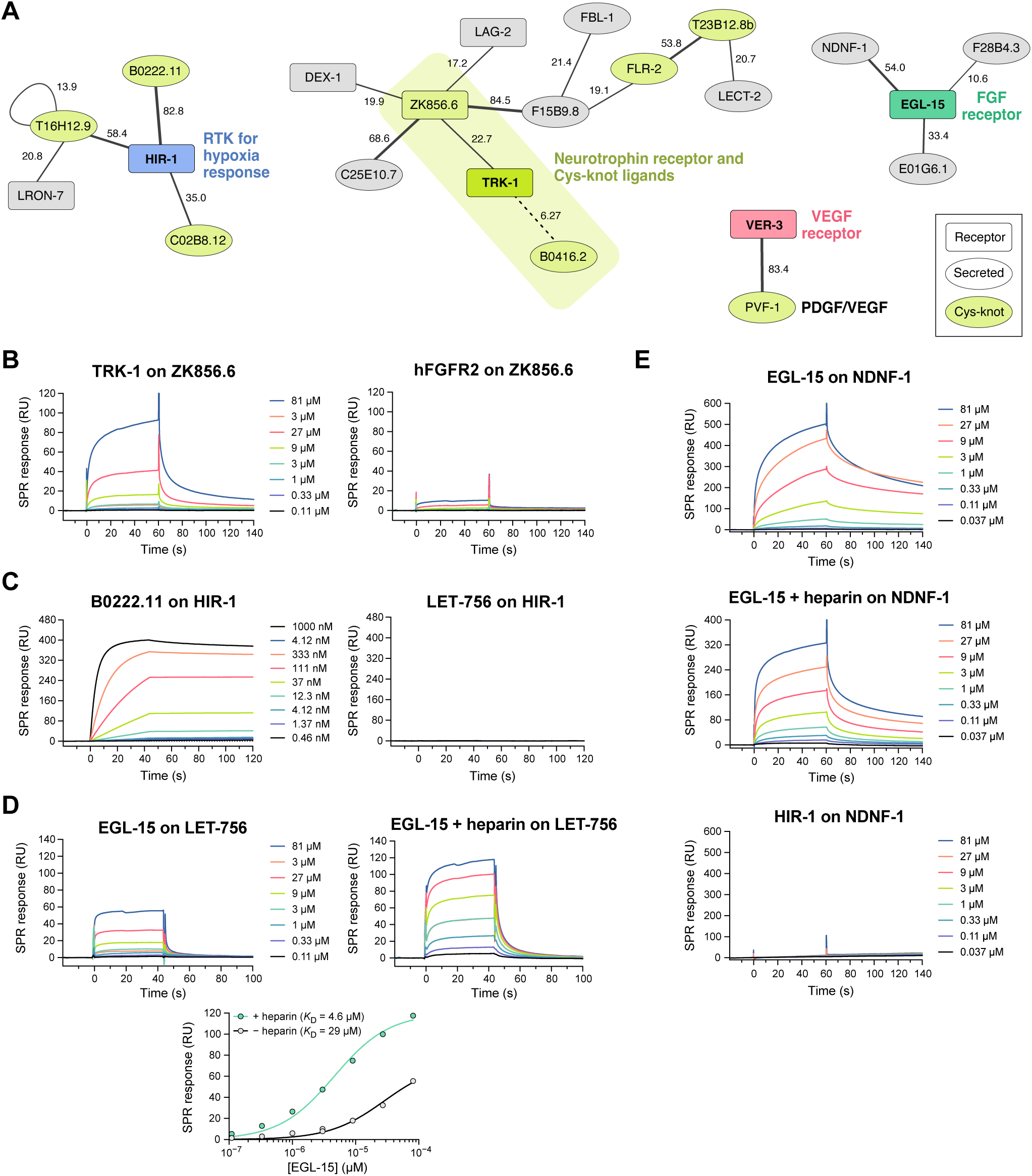
Networks of interactions with growth factor-like molecules and receptors. **A.** Network of interactions between cytokine- and growth factor-like molecules and receptors as observed in our screen (Figure 2). Line thickness corresponds to the symmetrized *z*-scores for each interaction. The numbers next to the lines indicate the *z*-score values. **B.** SPR sensorgrams for the interaction of TRK-1 with immobilized ZK856.6 and negative control (hFGFR2 against ZK856.6). **C.** SPR sensorgrams for the interaction of B0222.11 with immobilized HIR-1 and negative control (LET-756 against HIR-1). Kinetic fits and parameters are shown in Figure S5E. **D.** SPR sensorgrams and binding isotherms for the interaction of EGL-15 with immobilized LET-756 in the presence or absence of 50 μg/mL (3.1 μM) heparin. **E.** SPR sensorgrams for the interaction of EGL-15 with immobilized Fc-tagged NDNF-1 and negative control (HIR-1 against NDNF-1-Fc).

Among the family of Cys-knot family of secreted proteins, we observed that B0222.11, C02B8.12 and T16H12.9 interact with one receptor, HIR-1. This receptor was recently identified to direct hypoxia response and hypoxia-associated modeling of the extracellular matrix.^41^ HIR-1 is also a tyrosine kinase (RTK) with a cytoplasmic domain that resembles vertebrate RET and Fibroblast growth factor (FGF) receptors (FGFR). Based on this sequence similarity, the FGF homolog LET-756, which is a growth factor but not of the Cys-knot fold, was proposed as a potential HIR-1 ligand.^41^ Instead, we recommend that Cys-knot family proteins, including the three HIR-1 ligands, should be studied for hypoxia response in *C. elegans*. To validate our findings, we tested HIR-1 ectodomain against B0222.22 and LET-756/FGF with SPR using purified proteins (**Figure 5C**). We observed a high-affinity complex of HIR-1 with B0222.22 (*K*_D_ = 0.50 nM) (**Figure S5E**), while LET-756/FGF showed no binding to HIR-1.

We had included the two FGFs (EGL-17 and LET-756) and the FGFR ortholog (EGL-15) in our ectodomain collection. Surprisingly, we did not observe an interaction between these putative FGFs and the FGF receptor in our screen, likely as a result of complete lack of expression of the FGF ligands using our bait and prey plasmids in S2 cells (**Table S1**). To confirm this, we expressed and purified EGL-15/FGFR ectodomain and LET-756/FGF using lepidopteran cells and performed SPR experiments, which demonstrated the FGF-FGFR interaction in *C. elegans* (**Figure 5D**). This confirms that the lack of an FGF-FGFR hit in our screen is a false negative and was due to lack of protein expression. We observed stronger binding in the presence of heparin (**Figure 5D**), as previously observed for vertebrate FGFR-FGF interactions.^42^

In our screen, we observed NDNF-1, the nematode ortholog of the human neuron-derived neurotrophic factor, interacting with EGL-15/FGFR. NDNF and FGF are not homologous and do not share structural similarities, and an NDNF-FGFR interaction was not previously suspected or reported. However, NDNF overexpression is known to inhibit FGF signaling in cultured vertebrate cells, and FGFR-1 and NDNF are both implicated in congenital hypogonadotropic hypogonadism (CHH) and Kallmann syndrome (KS).^43,44^ Our discovery of an NDNF-FGFR complex, may provide the missing mechanistic link that connects NDNF with FGFR signaling, especially in disease states.

To validate the interaction of NDNF-1 with EGL-15/FGFR, we set out to express NDNF, but failed to produce stable protein with common expression systems. Therefore, we used Protein A-coupled SPR chips to capture Fc-tagged NDNF-1, originally produced as bait for ECIA, and measured binding to EGL-15 as an SPR analyte. We observed strong binding in the absence and presence of heparin (**Figure 5E**), demonstrating the validity of the NDNF-1–EGL-15/FGFR interaction.

Finally, *C. elegans* has a single platelet-derived growth factor (PDGF) homolog, PVF-1. In our screen, we observed that PVF-1 binds VER-3, the predicted PDGF/VEGF receptor ortholog, suggesting that the PDGF/VEGF-mediated biology is conserved from nematodes to mammals.

### *C. elegans* Wirins: ZIG-8 and RIG-5 and their interactions

An important result of our previous interactome study was the discovery of Dpr and DIP protein families (32 total members) in the fruit fly.^4^ Dprs and DIPs interact with each other, and have been strongly associated with synaptic specificity, axon guidance and fasciculation, cell fate determination and survival, and animal behavior.^45^ We recently identified the nematode homologues of Dprs and DIPs, ZIG-8 and RIG-5, respectively, as part of a conserved family named Wirins,^45^ and our interactome dataset reports a high-confidence ZIG-8–RIG-5 interaction as expected (**Table S3**). While homophilic and heterophilic interactions between Dprs and DIPs, and among their vertebrate orthologs (IgLONs) have been well established by others and us, it is unclear how Dprs and DIPs signal since they are GPI anchored proteins with no intracellular domains.^46^ No extracellular cell surface receptors have been reported as binding partners for Dprs and DIPs to-date.

In our dataset, we observed several interactions with ZIG-8/Dpr and RIG-5/DIP (**Figure 6A**). These include ZIG-8 binding to C34C6.3, and RIG-5 interactions with B0507.1, PTP-3, NLR-1, T19D12.6 and HMR-1. Among the binding partners, NLR-1, T19D12.6 and HMR-1 belong to the LamG+EGF family of proteins, most prominently represented by the synaptic Neurexin proteins. NLR-1 is the *C. elegans* ortholog of Contactin-associated proteins (CNTNAP), known as Nrx-IV in flies. Since CNTNAPs are known to take part in the formation of various types of cell junctions (such as *C. elegans* gap junctions, *Drosophila* septate junctions and vertebrate axo-glial junctions),^47–50^ and have known neuronal and synaptic developmental functions,^51,52^ they are plausible candidates for mediating Dpr/DIP signaling. The other strong candidate interaction for mediating Dpr/DIP signaling is the RIG-5–PTP-3 interaction: PTP-3 is a receptor protein tyrosine phosphatase (RPTP) and the worm ortholog for LAR. LAR has been implicated in *Drosophila* to control synaptic development and signaling in the photoreceptor cells and neuromuscular junctions,^53,54^ where Dpr-DIP was previously shown to control synaptic signaling.^55^ HMR-1 has been shown to be required for axon fasciculation and dendrite extension in *C. elegans*,^56–58^ and its fly ortholog, CadN, functions in synaptic targeting in the developing optic lobe;^59,60^ Dprs and DIPs are similarly known to control synaptic targeting of photoreceptor neurons.^55^ These strong functional links support the validity of these interactions. To further confirm these interactions, we performed SPR with purified RIG-5 ectodomain against PTP-3, NLR-1 and HMR-1 ectodomains, and validated the interaction hits (**Figures 6B-E**). These interactions proved to be lower affinity, as would be expected for co-receptors residing on the same cell, which may also require the formation of high-density ZIG-8/Dpr–RIG-5/DIP clusters.

**Figure 6.**
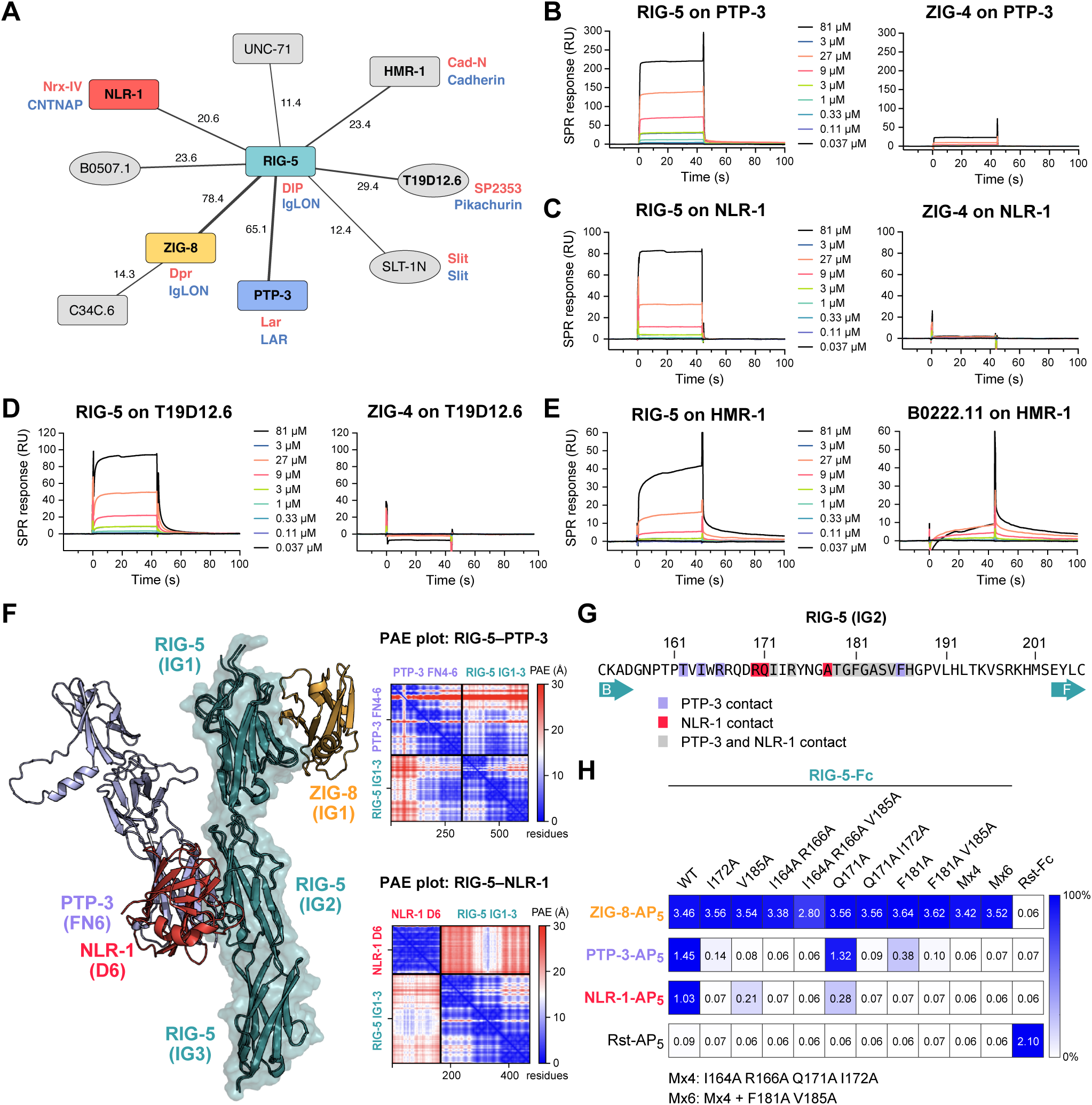
RIG-5 interacts with molecules with neuronal and synaptic functions. **A.** Interactions of ZIG-8/Dpr and RIG-5/DIP according to our screen. Red and blue names represent orthologs in *D. melanogaster* and mammals, respectively. Line thickness scales with the symmetrized *z*-scores for each interaction. The numbers next to the lines indicate the *z*-score values. **B-E.** SPR sensorgrams for interactions of RIG-5 with immobilized PTP-3 (B), NLR-1 (C), T19D12.6 (D), and HMR-1 (E). **F.** Overlay of the ZIG-8–RIG-5 structure (PDB ID: 6ON9) (Cheng *et al*., 2019) and AlphaFold-multimer models for RIG-5 IG1-3+NLR-1 D6 and RIG-5 IG1-3+PTP-3 FN4-6. ZIG-8/Dpr binding to RIG-5/DIP does not overlap with PTP-3 and NLR-1, which share a binding site on the RIG-5 IG2 domain. See Figure S6 for details of the AlphaFold-predicted interfaces. Predicted aligned error (PAE) plots for AlphaFold predictions are shown for both RIG-5 (ECD)-PTP-3 (FN4-6) and RIG-5 (ECD)-NLR-1 (D6) complexes. **G.** RIG-5 IG2 residues identified to interact with PTP-3 and NLR-1 in AlphaFold models. **H.** ECIA for RIG-5 ECD mutants against ZIG-8, PTP-3 and NLR-1 ECDs. Mutations in RIG-5 IG2 that break PTP-3 and NLR-1 binding do not affect ZIG-8 binding, which is known to happen through the IG1 domain. The raw readout for the assay (absorbance at λ = 650 nm) is noted in each square. The homodimeric Rst interaction is a positive control.

To gain further insights into some of these interactions, we used the Colabfold implementation of AlphaFold2-Multimer.^61,62^ We predicted models for both the RIG-5–PTP-3 and RIG-5–NLR-1 complexes with reasonable ipTM (interface predicted template modeling) values (0.72 and 0.49, respectively) and predicted aligned error (PAE) values (**Figure 6F**), where the second IG domain of RIG-5/DIP interacts with these signaling receptors using highly overlapping interfaces (**Figures 6F, 6G and S6**). We designed point mutations of RIG-5 that are likely to break these interactions, and tested them for binding via ECIA (**Figure 6H**): We observed that the mutations broke the PTP-3 and NLR-1 complexes of RIG-5 as predicted, but none affected ZIG-8/Dpr binding, which depends on an epitope on the first IG domain.^45,55^ These results support our RIG-5 interaction discovery, as well as the AlphaFold prediction of the binding interfaces. We suggest that the study of Dprs and DIPs in the fly model should include the orthologs Lar, Nrx-4 and CadN, to reveal Dpr-DIP signaling at the synapse.

### Limitations of the ECIA screen and further insights

While we observed many interactions that were previously known, there were some expected complexes that we did not observe. As mentioned above, the FGF-FGFR complexes were missed likely as a result of lack of FGF expression in our expression system. 23% of our constructs did not yield detectable expression, as judged by western blotting (**Table S1**), which is likely the largest source of false negatives. However, there are other unexpected negatives that cannot be explained by lack of expression based on known binding data with vertebrate homologs. One such case is the lack of binding between LAT-1 and TEN-1, the nematode homologs of Latrophilins and Teneurins, which form a synapse-instructive complex in mammals. A previous study suggested that this interaction may not exist in invertebrates on the basis of genetic data, and therefore the nematode and mammalian Latrophilins and Teneurins may act through different ligands.^63^ We report the Toll-like receptor TOL-1 and the LRR protein LRON-11 as interaction partners for LAT-1 and TEN-1 in *C. elegans*, respectively, and show in an accompanying paper that the LAT-1–TOL-1 complex is needed during early embryo development.^64^ Another unexpected observation from the high-throughput assay was the lack of an interaction hit between Neurexin (NRX-1) and Neuroligin (NLG-1). Neurexins and Neuroligins are major regulators of synapse formation and function, and interact strongly with each other.^65^ However, biochemical proof of a direct interaction between *C. elegans* NRX-1 and NLG-1 is limited.^66^ To scrutinize our unexpected negative result, we performed surface plasmon resonance experiments with the ectodomain of NLG-1 against the LNS6 domain of NRX-1, previously established as the domain responsible for Neuroligin interactions.^67^ Unlike studies demonstrating nM affinity with comparable constructs of mammalian homologs,^68,69^ we observed binding between nematode NRX-1 and NLG-1 with a ∼48 µM dissociation constant, about three orders of magnitude weaker than the mammalian orthologs (**Figures S7A and S7B**). Similarly, size-exclusion chromatography experiments showed no stable complex formation, unlike previous observations for mammalian Neurexin-Neuroligin complexes, but in agreement with a very weak complex (**Figure S7C**).^70^ Therefore, it is possible that while nematode Neurexins and Neuroligins may have preserved their neuronal functions, they may mediate it through interactions via other proteins, such as with MADD-4,^71^ via novel interaction partners we identified in our assay, such as TMEM132 for NRX-1, or heavily depend on other factors, such as heparan sulfate modifications of Neurexin as recently discovered^72^ for strong complex formation.

### AlphaFold-Multimer predictions for interactions observed by ECIA

Since we observed that AlphaFold models have proven useful in revealing novel interactions from our dataset, we took a systematic approach to study if AlphaFold results may indicate complex formation. We calculated AlphaFold-multimer complexes for 72 high-confidence interactions we observed for complexes with <2500 amino acids, as well as 46 complexes created by randomly pairing proteins from the same set of proteins. We used the ipTM values reported by AlphaFold as a measure of its confidence for the complex model. Our observed complexes had an average ipTM value of 0.495 ± 0.241, while random protein pairs gave an average ipTM value of 0.273 ± 0.139 (**Figure 7A and Table S3**). The remarkable differences in the distribution of ipTM values for observed complexes vs. randomly paired proteins attest to the usefulness of AlphaFold as a predictor of protein complex formation, while also supporting our discovery results.

**Figure 7.**
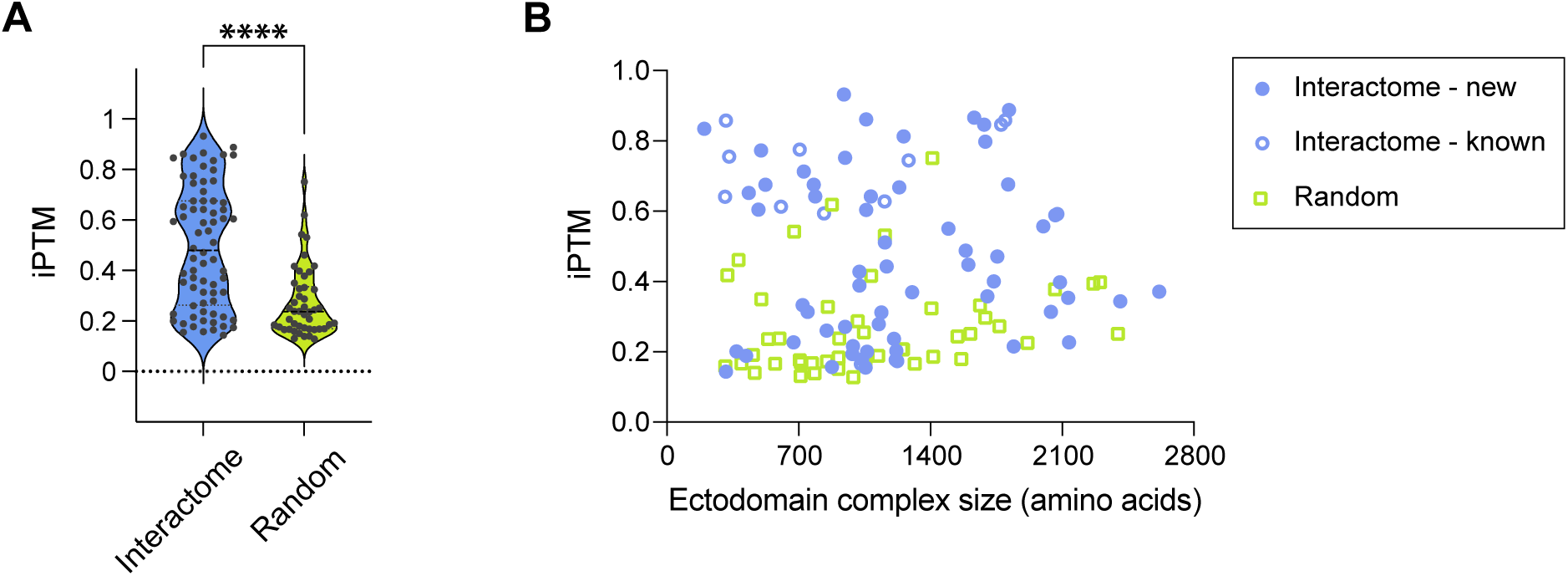
AlphaFold predictions of extracellular protein complexes and chemical synapses corresponding to PPIs. **A.** ipTM (interface predicted Template Modelling) score values of AlphaFold-predicted complexes from our interactome (blue) and random pairings of proteins (green). **B.** Size of the protein complex does not correlate with ipTM values. ECIA hits with previously determined homologous structures (open blue circles) have the highest ipTM values on average.

To analyze AlphaFold predictions further, we investigated whether observing higher ipTM values correlate with any property of our complexes. We observed no correlation between sequence length and ipTM (**Figure 7B**). However, ipTM values were significantly higher for complexes which had homologous (mostly mammalian) complex structures in the Protein Data Bank (0.676 ± 0.190). This is not surprising, as these structures were likely in the training set for AlphaFold-multimer. However, even after removing protein pairs with homologous complex structures in the PDB, the complexes we detected had significantly higher ipTM values on average (0.459 ± 0.236) than randomly paired complexes, demonstrating that AlphaFold can be a useful tool in identifying novel protein complexes, and for studying the structures of these complexes. Computational pipelines using this idea have been established recently.^73^

### Chemical synapses corresponding to the observed interactions

Protein-protein interactions (PPIs) play a crucial role in forming chemical synapses between neurons.^74^ Therefore, we investigated the chemical synapse connectome^75^ associated with PPIs. Specifically, a PPI is considered as evidence that supports a chemical synapse if the corresponding genes are expressed in the neurons forming a chemical synapse. We used the gene expression data from CeNGEN.^76^ Among the 134 PPIs that has available gene expression data related to chemical synapses, there are only 4 PPIs that do not serve as evidence of any known chemical synapses, namely ZC374.2–T02C5.1A, INS-6–ZIG-2, SGCA-1–IRLD-51, and LRON-7–T02C5.1A. The remaining 130 PPIs can potentially explain the entire known chemical synapse connectome. While 99.9% of chemical synapses have more than 5 evidences, there are a few examples of more specific chemical synapses. For example, HMR-1–HMR-1, IGCM-3– IGCM-3, and MAB-20–PLX-2 support the chemical synapses between the AVKL and AVKR neurons. Additionally, we have included the 185 experimental PPIs with the number of supporting synapses in **Table S7**. To establish a robust statistical framework for evaluating the significance of these interactions associated with the formation of chemical synapses, we implemented a randomized neuron connectome as a control. Our analysis revealed two distinct categories of noteworthy PPIs: (1) PPIs with the highest number of supporting synapses: TMEM-132–NRX-1, CASY-1–T19D12.6, PTP-3–NLR-1, CASY-1–PLX-1, SAX-3–PLX-1. These interactions are of particular interest due to their prevalence within the synaptic connectome, potentially indicating their functional importance in neuronal communication; (2) PPIs demonstrating the highest statistical significance when compared to the randomized control: SLT-1N–SLT-1C, LECT-2– MIG-13, UNC-5–PXN-1, ZIG-4–DAF-28, F15B9.8–ZK856.6. The statistical significance of these PPIs suggests that their occurrence is non-random and may represent biologically relevant interactions worthy of further investigation. These results highlight significant putative contributions of experimental PPIs in unraveling the intricate complexities of chemical synapses.

## DISCUSSION

Protein-protein interactions in the extracellular space control the development and functioning of all aspects of multicellular physiology. With this study, we report the results of an extracellular interaction screen of 379 *C. elegans* cell surface receptors and secreted proteins, the largest for a non-human species to-date, that covers a diverse range of the protein fold space. The interactions we report are mostly novel, even when counting those identified for orthologs in other taxa as previously known (86% previously unknown).

### Novel interactions among guidance cues and receptors for neuronal development and animal morphogenesis

In our ectodomain collection, we have included known axon guidance cues and receptors, and to our surprise, observed many novel interactions that link separate cue-receptor axes. This brings up the possibility that these interactions might build “supra-signaling complexes” able to incorporate cues and signaling receptors from multiple pathways. Based on our map of domain-domain interactions (**Figure 3H**), the Robo-Ephrin-Semaphorin (SAX-3–EFN-4–MAB-20), Robo-Ephrin-Eph (SAX-3–EFN-4–VAB-1) or Robo-Plexin-Semaphorin (SAX-3–PLX-1–SMP-1/2) supercomplexes may be possible, as the domains needed to form these complexes do not overlap. How these supra-molecular complexes would signal needs to be studied *in vivo*, especially in the light of expression patterns of these various proteins. It is also possible that some of the new interactions we identified act to silence the canonical pathways by blocking the formation of the known complexes: this may be the case for SAX-3/Robo, as all its binding partners interact with its IG domains. These interactions can be studied readily, thanks to the extensive collection of mutant strains, published genetic interactions, and expression data already available in *C. elegans*.

The interactions we identified between the separate axon guidance axes also highlight the multiple roles these receptors and ligands play. For example, morphogenesis of the intestine in *A. elegans* is known to be mediated by movements of cells governed by the actions of MAB-20/Semaphorin, SAX-3/Robo and EFN-4/Ephrin, which we have now identified to interact with each other.^77^ Similarly, *efn-4* and *mab-20* mutant animals have highly similar phenotypes in axonal growth and guidance in the same cells,^78,79^ as well as in epidermal enclosure of embryos,^80^ and male tail morphogenesis.^81,82^ Genetic interactions and shared phenotypes identified for these genes are likely due to the direct physical interactions we report here.

### Growth factors and their receptors in *C. elegans*

We have identified cystine-knot binding partners for the *C. elegans* Trk-like receptor, TRK-1. Previous efforts to identify nematode neurotrophins and their receptors by sequence have not been successful.^83^ Despite lack of apparent sequence similarity for both the vertebrate neurotrophins and the Trk receptor ectodomain, AlphaFold prediction of our newly identified complexes at 2:2 stoichiometry, show strong resemblances to the known structures of mammalian Trk receptors, neurotrophins and their complexes (**Figure S4**).^84–86^

We report additional interactions for growth factor-like molecules, one of which is the worm ortholog of the Neuron-derived neurotrophic factor, NDNF-1, interacting with EGL-15/FGFR. We suggest that this interaction may be the underlying reason for why NDNF and FGFR mutations are both found in human patients with congenital hypogonadotropic hypogonadism and Kallmann syndrome. The fruit fly NDNF, Nord, also binds with and regulates degradation of Dally, a glypican-type heparan sulfate proteoglycan.^87,88^ Since FGF-FGFR interactions are strengthened by heparan sulfate (**Figure 5D**),^56,57^ and that heparin does not break the NDNF-1-EGL-15/FGFR interaction (**Figure 5E**), it is reasonable to speculate that NDNF may interact with Dally/Glypican as well as FGF receptors, and prevent FGF from turning on FGFR signaling.

### Interactomes for model organisms

One important takeaway message we learned is that proper study of biological processes in a model organism requires dedicated effort in revealing the relevant biochemistry for that model organism. As one example, insulin signaling in *C. elegans* has been heavily studied and proven insightful for understanding aging, neuronal and embryonic development. However, as we show here, more may need to be done to reveal the biochemical components of the system (such as ZIG-2 to -5) and their effects on signaling, which may or may not be shared with other model organisms.

Conversely, our interactome dataset will likely prove useful across many model organisms. For example, we revealed several new binding partners for the two *C. elegans* homologs of *Drosophila* Dprs and DIPs, which are a 32-member receptor family known for their roles in neuronal wiring in the fruit fly. Our findings will facilitate study of neuronal wiring in both model systems, given the novel signaling co-receptors we propose for this receptor family. In addition, the interactome dataset can be used in conjunction with expression data to identify candidate proteins and complexes with roles in any cell surface process in specific cell types, such as synapse formation. For example, expression of ZIG-8/Dpr and RIG-5/DIP are enriched in pairs of cells known to form synapses (**Table S7**).

### Towards a complete extracellular interactome

Here, we report significant improvements to our extracellular interactome methodology, including a statistically rigorous method for data analysis, applicable to related interactome strategies and high-throughput ELISA-like assays. We also implemented enhancements to improve throughput; Additional gains in throughput can also be achieved by pooling of bait or prey, as recently done in Wojtowicz *et al.*^8^ The largest obstacle for us has been the time and cost of creating the plasmid collection, while the entire assay could then be completed by two full-time researchers in only two months. As gene synthesis costs decrease significantly, complete interactomes of all receptors and secreted proteins will become cheaper to execute for several model organisms of interest, to support a more complete mechanistic understanding of development, health and disease.

## METHODS

### Selection of Ectodomains

We collected names of genes annotated to have selected domains from *SMART*^89^ and *SUPERFAMILY*^90^ databases in the *C. elegans* genome. Transcripts for these genes were identified in the WS252 release of the *C. elegans* genome, and analyzed using *phobius* for signal peptide and transmembrane domains.^91^ The identified ectodomains were written into Genbank-formatted files for bait and prey plasmids for cloning and/or gene synthesis. The bioinformatic pipeline described above was performed using tools available in Bioperl modules. Proteins that have no discernable transmembrane domains were also run on the PredGPI^92^ server (http://gpcr.biocomp.unibo.it/predgpi/) to assess whether they may be GPI-anchored; we removed the C-terminal GPI anchoring sequences for those proteins with predicted GPI-anchors. Predictions were repeated with the recently released NetGPI^93^ to update **Table S1**.

Following curation of ectodomain sequences, included in **Table S1**, we re-analyzed domain compositions using updated tools and AlphaFold predictions,^94^ which became available as we prepared the manuscript for publication. Based on the results from these tools, we decided not to distinguish EGF domains from WR1 domains (SMART SM00289; Interpro IPR006150), EB modules (Pfam PF01683) and Lustrin-type Cys-rich domains (Pfam PF14625; Interpro IPR028150), which were previously defined using sequence alignments. We observed that domain prediction tools such as *SMART* and *hmmscan*^95^ repeatedly gave overlapping predictions for these families, and structure predictions showed strong similarities and a continuum of elaborations above basic features, making a rigid distinction between these folds difficult. Future work using experimentally determined structures, predictions and sequence alignments will be necessary to carefully classify similar Cys-rich small domains into better-defined categories.

### Ectodomain Expression Library Construction and Protein Expression

To facilitate protein expression, we built new ECIA vectors carrying the strong constitutively active Actin 5C promoter rather than the inducible metallothionein promoter on pECIA2 and pECIA14 described in Özkan *et al*.^4^ Briefly, the multiple cloning site (MCS) region of pACTIN-SV^17^ was replaced by a BiP signal sequence followed by the cassette attB1-MCS-attB2-3C-Fc-V5 tag-His_6_ (3C: HRV 3C Protease site) to generate the bait vector pECIA75, while the replacement of the MCS of pACTIN-SV by BiP-attB1-MCS-attB2-3C-COMP-AP-FLAG-His6 (COMP: pentameric coiled coil from rat COMP; AP: human placental alkaline phosphatase) resulted in the prey vector pECIA76.

To construct the expression library for the proteins of interest listed in Table S1, the ectodomains of 16 proteins from our lab collection were subcloned into the MCS of the new ECIA vectors. We also purchased a *C. elegans* ORF library from GE Healthcare and found 107 proteins in Table S1 from the 12611 clones of the ORF library. However, only 63 (including 5 with mutation(s) or intron(s) that were corrected in cloning) were successfully subcloned while the remaining 44 clones from the clone collection proved to be wrong or empty vectors. Through RT-PCR, 9 were amplified from a *C.elegans* N2 mRNA pool (gift from Kang Shen at Stanford). The left 288 were synthesized by GenScript.

Every bait or prey construct was individually expressed using transient transfection of *Drosophila* Schneider 2 (S2) cells in Schneider’s medium with 1.8 g/L L-Glutamine (Gibco, 21720024) with 10% Fetal Bovine Serum, 50 units/ml Penicillin, and 50 μg/ml Streptomycin. Briefly, 10 mL of S2 cells at 1.8 million per mL were seeded in T75 flasks and incubated at 28°C overnight. The cells were transfected transiently next day with 5 μg of each plasmid using TransIT®-Insect Transfection Reagent (Mirus, MIR 6104) following manufacturer’s manuals. Conditioned media were collected 4 days after transfection. Protease inhibitors (Sigma, P8849) and 0.02% NaN_3_ were added to harvested media before storage at 4°C in 15 mL conical tubes.

All bait and prey samples collected were run on SDS-PAGE gels, blotted and probed with mouse THE™ His Tag Antibody [iFluor 488] (GenScript, A01800) for assessing protein expression. Bio-Rad ChemiDoc Imaging System was used to quantitate the bait and prey samples based on the reference band of 0.1 µM rat His_6_-tagged GluD2 ectodomain (ATD+LBD)^96^ on each blot. Overall, we observed expression as high as ∼0.31 μM (LRON-3-Fc) in conditioned media. The lowest protein concentration we could detect and measure was ∼0.45 nM, for R09H10.5-AP_5_. Expression for 23% of the samples could not be detected. Relative expression levels are reported in **Table S1**.

### Extracellular Interactome Assay (ECIA)

Clear Nunc 384-well MaxiSorp plates (Thermo Scientific, 464718) were coated with 20 µl 5 µg/mL Protein A (Genscript, Z02201) in 100 mM sodium bicarbonate pH 9.6 at 4°C overnight. Excess protein A was discarded, and the coated plates were blocked with 90 µl 10% SuperBlock™ T20 (PBS) Blocking Buffer (ThermoFisher, 37516) in PBS at 500x rpm for 3 hours at room temperature. The blocking buffer was then removed, and the plates were washed three times with 90 µl PBST (PBS + 0.1% Tween 20) using a microplate washer (BioTek). 20 μl of medium containing a secreted Fc-fusion protein (bait) was added into wells of a single plate at 4°C with a shaking at 500 rpm on a plate shaker overnight for bait capture. Next day, plates were blocked with 1% Bovine Serum Albumin (BSA) in PBS for three hours at room temperature while shaking at 500 rpm, followed by three washes with 90 µl of wash buffer (PBS with 1 mM CaCl_2_, 1 mM MgCl_2_, and 0.1% BSA). This was followed by addition of 20 μl of different medium containing secreted AP_5_-fusion proteins (prey) into each well of a plate using a Rainin BenchSmart 96-well multipipetter and incubation at room temperature while shaking at 500 rpm for 3 hours. The plates were then washed three times with 90 µl of wash buffer. Finally, 50 μl of BluePhos Phosphatase Substrate (KPL, 50-88-02) was added to each well using a Rainin BenchSmart 96-well multipipetter at room temperature. Absorbance at 650 nm was measured at 1 and 2 hours using a microplate reader and images of these 384-well plates were scanned.

### The contents of the large high-throughput screen data

Each 384-well plate included four control measurements and 379 prey paired against one bait (+ one prey, NAS-23, repeated as a control but not used in the analysis). The 379 unique ectodomains correspond to 374 genes, where some genes were represented by more than one constructs, such as the N- and C-terminal fragments of SLT-1 (see Table S1).

The large ECIA screen data has each pairwise heterotypic interaction tested twice, with each protein alternatively in the bait vs. prey configuration; this yielded two separate reports of interaction strength. Homotypic interactions were measured once.

### Analysis of ECIA Data: MaxEnt statistical model

To assess the significance of interactions, we employ a maximum entropy network ensemble-based technique^19^ to construct a statistical model that captures the absorbance background. Initially, we represent the experimental network by an *n* × *m* matrix *A* and normalize it as *An* = *A/A_max_* (**Figure S2A**). Here, the rows and columns denote prey and bait, respectively, while entries denote the normalized absorbance of each candidate protein-protein interaction (PPI). To capture the random background given the observed row and column sums of *An*, we maximize the entropy of a random network ensemble, denoted as *S* = −*Σ_G_P*(*G*) ln *P*(*G*), subject to soft constraints that preserve the average row and column sums of the normalized experimental data *An*. Here, *G* denotes random networks and *P* denotes the probability of *G*. With maximized entropy, the model generates an ensemble with the broadest distribution of binary networks (a matrix with only 0 and 1 entries) that on average replicates the row and column sums of *An*. The average matrix of the *P* serves as the baseline expectation with the absence of any biological information (Figure 1B). Each matrix entry in *P* representing the connection probability of a prey-bait pair as *p_ij_* = 1/(*α_i_β_j_* + 1). In practice, the parameters *α_i_*,*β_j_* are iteratively determined to preserve the mean total normalized absorbance of each prey *k_i_* = Σ*_j_α_ij_* and bait *k_j_* = Σ*_i_α_ij_* as

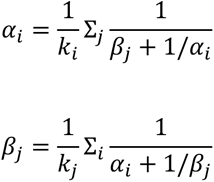

where *α_ij_* represents the elements in the normalized absorbance matrix *An*. Here we apply 100 iterations to ensure the convergence of *α_i_* and *β_j_*. With these optimized *α_i_* (*β_j_*) for each row (column), we test the self-consistency of the model by verifying the implication that all *α_ij_* should satisfy *α_ij_* = 1/(1 + *α_i_β_j_*). However, upon using the optimized *α_i_* and *β*_*j*_, we obtain sets of *m* (*n*) different values of *α_i_* (*β_j_*) for each *α_i_* (*β_j_*), calculated as 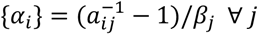 and 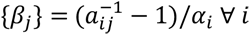. We observed small fluctuations of these {*α_i_*} and {*β_j_*} around the optimized values, quantified as 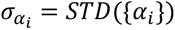 and 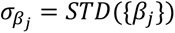. The standard deviation of *p_ij_* is then calculated using error propagation as 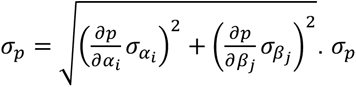 quantifies the systematic error that emerges from the deviation of the experimental data from the statistical model. In principle, there is also statistical error in *p_ij_* that can be calculated by 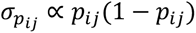. However, we find that the systematic error dominates in this case, thus, the statistical error is ignored. *z*-scores are used to identify potential signals that are significantly different from the baseline expectation as 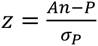.

Although the *z*-score matrix can be asymmetric, we anticipate that a large fraction of the PPIs are symmetric, meaning reciprocated signal in the two experimental directions (*s* → *t* or *t* → *s*). Thus, the reciprocal ratio of interactions works as a metric to reflect the precision of the experiment, akin to a cross-validation scenario. In **Figures S2D-E**, we illustrate how the reciprocal ratio of interactions varies with different z-score thresholds and unique PPIs. We assign z-score to each PPI by symmetrizing z scores in both directions as *z* = (*z_A_*_→*B*_ + *z_B_*_→*A*_)/√2 (**Table S2**). The scaling factor √2 ensures that the combined z-score remains a standard z-score, allowing for direct comparison and interpretation. We use the reciprocal ratio to determine the ideal z-score threshold to distinguish signal from background noise. Our analysis demonstrates that the maximum reciprocal ratio of 0.67, corresponding to 145 unique PPIs, is achieved at a stringent threshold of *z* = 12.2. Additionally, we observe that the reciprocal ratio fluctuates slightly between 0.61 and 0.67 for z-scores between 8.4 and 33.6 (**Figure S2D**). As a result, we propose an intermediate threshold of *z* = 8.4, which includes 185 unique PPIs (**Figure S2C, Table S3**). Selecting an even lower threshold is expected to include more false positives as indicated by a diminishing reciprocal ratio.

### Network Analysis of ECIA Data and Communities

Network theory offers a particularly powerful framework for mining insight from measurements of physical protein-protein interactions. Here, we conducted a network analysis of our ECIA dataset.

*Community detection:* We identified communities in this PPI network using a multi-scale partition algorithm,^21^ which identifies node groupings of different sizes by random walks of various lengths. The scale of the communities obtained by this approach is controlled by a hyperparameter tau (resolution), analogous to the length of the walk. Smaller values of tau correspond to shorter walks, which will on average end relatively close to where they begin. This results in the identification of relatively small communities. As the length of the walk (tau) increases, increasingly large neighborhoods are reached, with the likelihood of remaining in any particular neighborhood of the network as the walk proceeds determined by how insular the connectivity is within that neighborhood. We used this method across 50-fold variation in tau (0.1 to 5) to compute community partitions at a variety of scales. For each protein, we recorded the union of all other nodes that appear in the same community, across all choices of tau. The subset of proteins grouped in the same community across at least 25% of the tau range and *z*_min_ range are further labeled as “canonical” community neighbors, and are made available in Table S4.

Network analyses were performed using custom code written in Python 3.9, using NetworkX and python-louvain (https://github.com/taynaud/python-louvain). Network visualization was performed using NetworkX and Gephi. Code, data and output neighbor lists are available at https://github.com/mattrosen/ECIA-network-analysis.

### Protein Expression and Purification for Interaction Validation

For crystallography and SPR experiments, proteins were produced using baculoviral infection of *Trichoplusia ni* cells cultured in Insect-XPRESS (Lonza, BP12-730Q) or ESF 921 (Expression Systems, 96-001) media. Ectodomains or smaller fragments of proteins were cloned into baculoviral transfer vector pAcGP67 and baculoviruses viruses were generated with BestBac linearized DNA (Expression systems, 91-002) using Sf9 cells cultured in Sf-900 III (Gibco, 12658019) with 10% FBS. The constructs were cloned with the addition of C-terminal biotin acceptor peptide (for biotin capture on strepatividin-based SPR chips) followed by a hexahistidine tag. The constructs used in crystallography were cloned with C-terminal hexahistidine tags only. Proteins were purified from insect cell media using affinity chromatography with Ni-NTA agarose resin (Pierce HisPur, 88223) followed by size-exclusion chromatography using Superdex 200 Increase or Superose 6 Increase 10/300 GL columns (Cytiva) in 10 mM HEPES, pH 7.2 or 7.4, 100 mM NaCl (HEPES-buffered saline; HBS). Proteins used in SPR experiments were usually biotinylated using BirA ligase (Avidity, BirA500) and purified on size-exclusion columns to remove free biotin.

### Surface Plasmon Resonance Experiments

SPR experiments were carried out using SA or CM5 sensor chips with Biacore T200 or 8K models (Cytiva). Proteins were immobilized in HBSp+ (10 mM HEPES, 150 mM NaCl, 0.05% Tween 20), pH 7.2 or 7.4 using biotin capture on SA chips, or random amine coupling on CM5 chips when proteins were not tagged with a biotinylation sequence. Experiments were performed at 25°C, at a flow rate of 30 µl/min, with varying ligand immobilization levels (between ∼ 500 RUs to 1200 RUs), association and dissociation times, running buffer, and regeneration conditions. Details of the SPR experiments are listed in the **Table S6**. SPR sensorgrams and isotherms were plotted using Prism version 10.

For NDNF-1–EGL-15 SPR experiments (Figure 5E), 60 mL of S2 culture (in Schneider’s medium supplemented with Insect media supplement, Sigma I7267, and no serum) was transiently transfected with the Fc-tagged NDNF-1 (bait) expression plasmid, and conditioned medium was collected three days post-transfection. Medium containing NDNF-1-Fc was dialyzed overnight against HBS, pH 7.4, and was captured on Protein A SPR chips (Cytiva, 29127555) to validate the NDNF-EGL-15 interaction, using purified EGL-15 ectodomain as analyte. We observed slow dissociation of the NDNF-1-Fc from the SPR chip over several hours, resulting in a slow but noticeable baseline drift.

Except in cases where kinetics were too slow, we used the 1:1 Langmuir binding model for fitting binding isotherms to acquire *K*_D_ values in Prism version 10. With slow kinetics, we fit sensorgrams to kinetic models using Biacore’s BIAEvaluation software to acquire rate constants; we indicated which kinetic model was used in the relevant figure legends.

### Structure Determination for the ZIG-4–INS-6 complex

The ZIG-4–INS-6 complex was formed by co-expression using baculoviral expression of secreted ZIG-4 and INS-6. Both proteins were C-terminally tagged with hexahistidine tags, which allowed us to purify the complex from conditioned media using Ni-NTA Agarose resin. Complex was further purified over a Superdex 200 10/300 Increase (Cytiva) size-exclusion chromatography column in HBS. Purified complex was concentrated to 5 mg/ml, and screened for crystallization using the sitting-drop vapor-diffusion method at 22°C.

The ZIG-4–INS-6 complex crystallized in several conditions, which resulted in crystal structures resolved in three space groups. The tetragonal crystal form was grown in 0.1 M NaCl, 0.1 M sodium cacodylate, pH 6.5, 2 M (NH_4_)_2_SO_4_. These crystals were cryo-protected in a solution containing 0.1 M NaCl, 0.1 M sodium cacodylate, pH 6.4, 1.6 M (NH_4_)_2_SO_4_ and 24% glycerol, and diffracted to ∼1.3 Å resolution. We also grew crystals The *C*-centered monoclinic crystals were grown in 0.2 M sodium/potassium phosphate, 0.1 M bis-tris propane, pH 6.8, 28% PEG 3350. These crystals were cryoprotected in 0.2 M sodium/potassium phosphate, 0.1 M bis-tris propane, pH 6.8, 28% PEG 3350, and 20% glycerol, and diffracted to ∼2.3 Å resolution. Finally, the primitive monoclinic crystals were grown in 0.2 M sodium iodide, 0.1 M bis-tris propane, pH 6.5, 20% PEG 3350. These crystals were cryoprotected in 0.2 M sodium citrate, 0.1 M bis-tris propane, pH 6.5, 20% PEG 3350, and 25% glycerol, and diffracted to ∼2.4 Å resolution.

All crystallographic data were indexed, merged, and scaled using the *XDS* package.^97^ Molecular replacement was performed with *PHASER*^98^ in the *PHENIX* package^99^ using PDB ID: 6FEY (ImpL2 + DILP5) as the molecular replacement model^36^ with the tetragonal dataset. The refined higher-resolution model was used for molecular replacement with the other two datasets. *phenix.refine* was used to refine all models in reciprocal space and water placement,^100^ and *COOT* was used for model building and corrections in real space.^101^ Model building and refinement was guided by *MOLPROBITY* chemical validation tools within *PHENIX*.^102^

The coordinates and structure factors for the ZIG-4–INS-6 complex are deposited at the Protein Data Bank with the following accession codes: 8TK9 (tetragonal form), 8TKT (C-centered monoclinic form), 8TKU (primitive monoclinic form). Crystallographic data collection and model refinement statistics are in **Table S5**.

## ACKNOWLEDGMENTS

We would like to thank Kang Shen for guidance and sharing reagents, and Jingxian Li, Demet Araç, and Joseph S. Pak for technical and critical discussions, and Mateusz Krzyscik and Dengke Ma for sharing reagents. We acknowledge Hyun Lee and the University of Illinois at Chicago Biophysics Core Facility, and Elena Solomaha and the UChicago Biophysics Core Facility for SPR access and help. E.Ö. and I.A.K. acknowledge both support from the National Institute for Theory and Mathematics in Biology through the National Science Foundation (grant number DMS-2235451) and the Simons Foundation (grant number MP-TMPS-00005320). This study used resources of the Advanced Photon Source (APS), a U.S. Department of Energy (DOE) Office of Science User Facility operated for the DOE Office of Science by Argonne National Laboratory under Contract No. DE-AC02-06CH11357. Crystallographic data was collected at Northeastern Collaborative Access Team beamlines, which are funded by the National Institute of General Medical Sciences from the NIH (P30 GM124165). The Eiger 16M detector on the 24-ID-E beam line is funded by a NIH-ORIP HEI grant (S10OD021527).

## SUPPLEMENTAL INFORMATION

**Figure S1.**
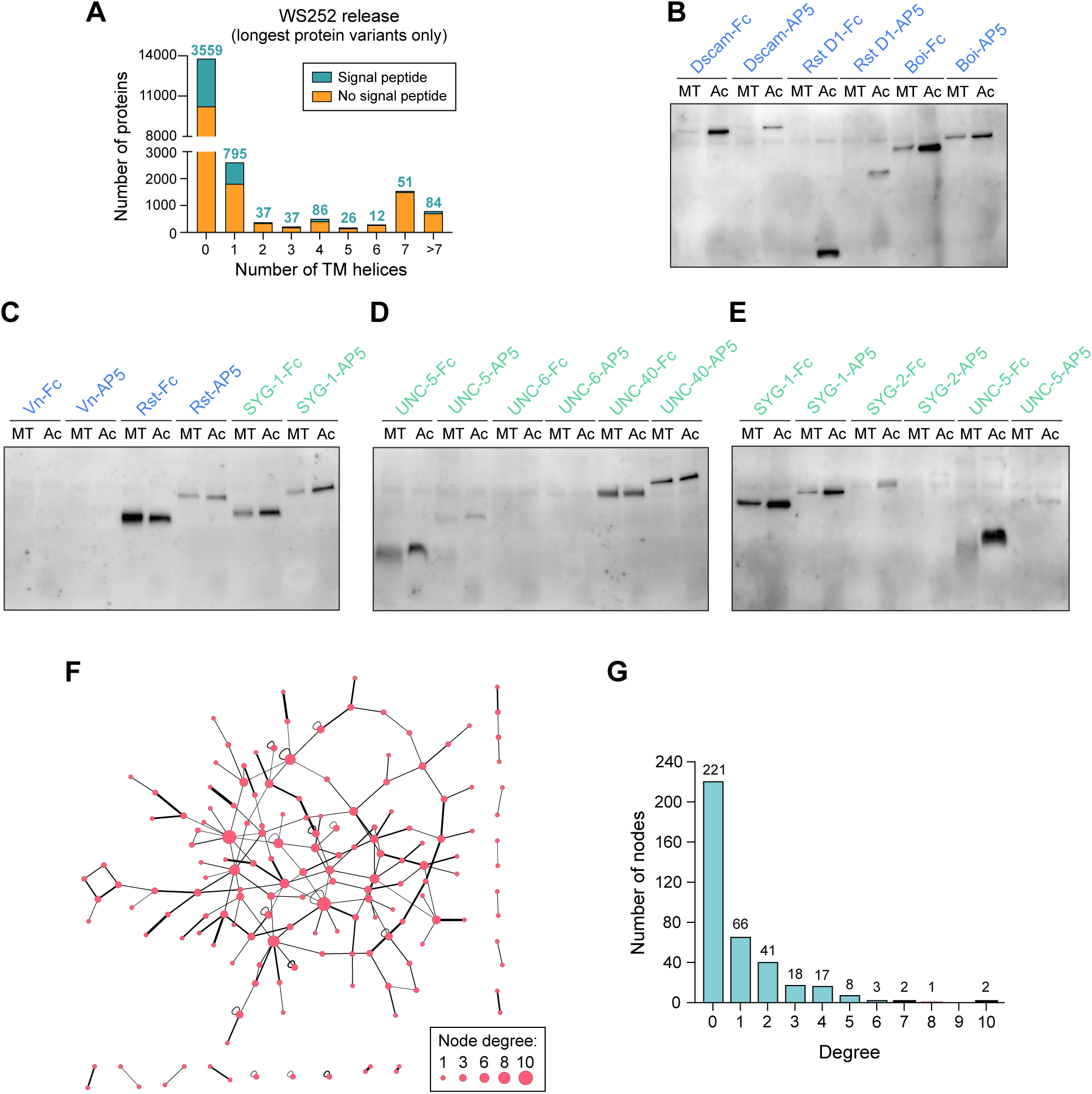
Domain collection, protein expression and network analysis. **A.** Signal peptide and transmembrane helix analysis of the *C. elegans* proteome (WS252 release) shows that 23% of proteins have predicted signal peptides, and 44% of proteins are either membrane-anchored or secreted. **B-E.** Expression testing of *D. melanogaster* (blue) and *C. elegans* (green) ectodomains in S2 cells using the Metallothionein (MT) and Actin 5C (Ac) promoters. Rst D1 refers to the fist immunoglobulin domain of Rst. For MT-driven expression, transiently transfected cells were induced with 0.8 mM CuSO_4_ at 16 hours post-transfection. All transfections were collected 88 hours post-transfection for western blotting using a mouse primary anti-His antibody (1:2000) and an Alexa Fluor 488-coupled donkey anti-mouse IgG secondary antibody (1:5000). Overall, the Actin 5C promoter results in higher expression, but not in every case. **F.** Network of 185 interactions detected with a cutoff of *z*_min_ > 8.4 drawn using the organic layout algorithm in Cytoscape, where node size relates to node degree (see the legend), and the edge thickness scales to *z*_min_. **G.** The degree distribution of all the interactions depicted in F.

**Figure S2.**
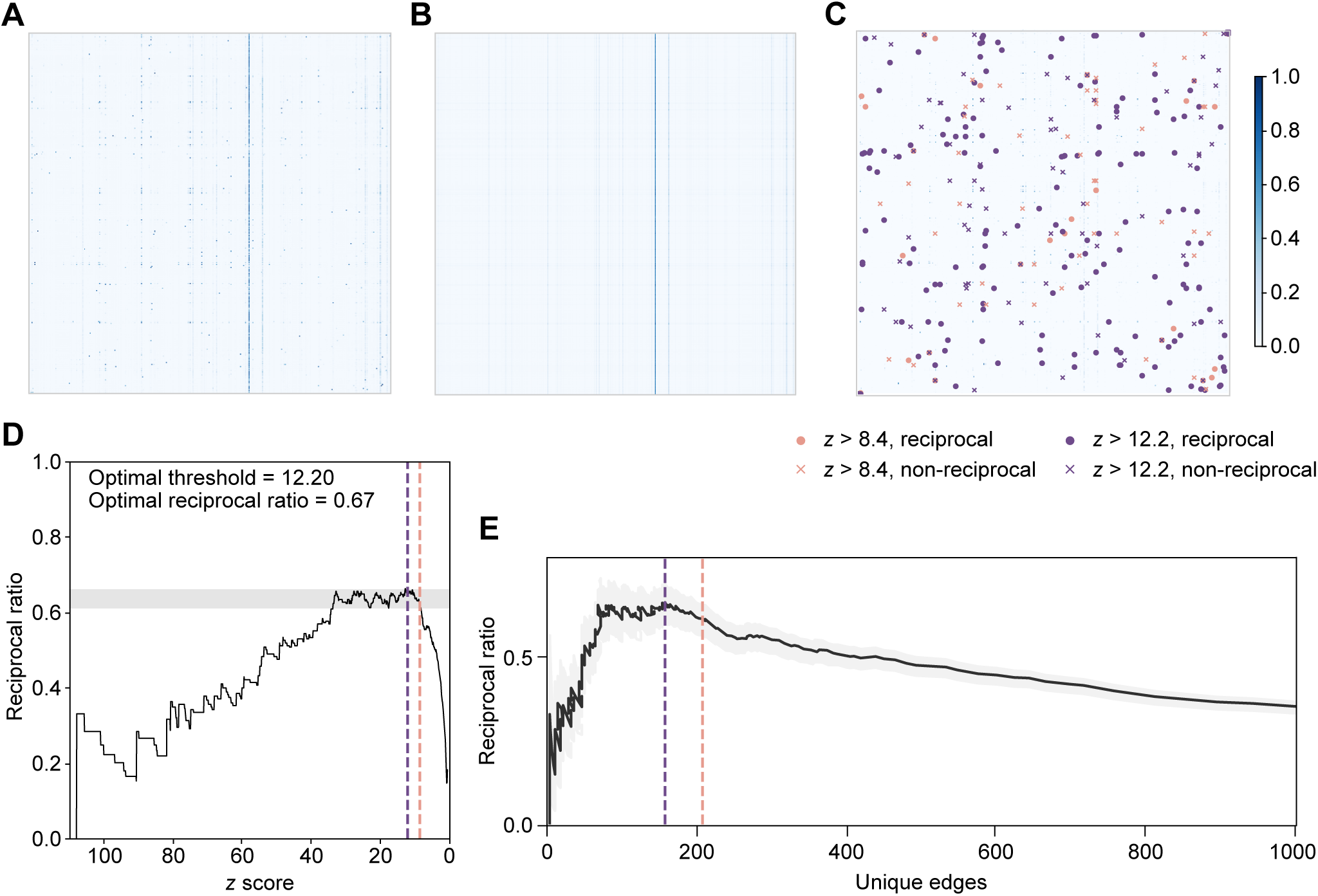
MaxEnt model to filter the experimental data. **A.** The normalized experimental data *An*. **B.** The mean of the statistical background model *P*. **C.** The difference between *An* and *P*. PPIs with z-score above intermediate (orange) and stringent (purple) thresholds are shown in matrix form. Reciprocal PPIs are marked with dots (**•**) and non-reciprocal PPIs are marked with an ‘×’. **D.** The reciprocal ratio of interactions as a function of the chosen threshold of *z*-scores. The maximum reciprocal ratio is achieved with *z* = 12.2. **E.** The reciprocal ratio as a function of the number of unique edges identified. The shading represents *n* ± *SE*, where *n* is the number of reciprocal edges. SE is calculated by the shot noise as *SE* = √*n*.

**Figure S3.**
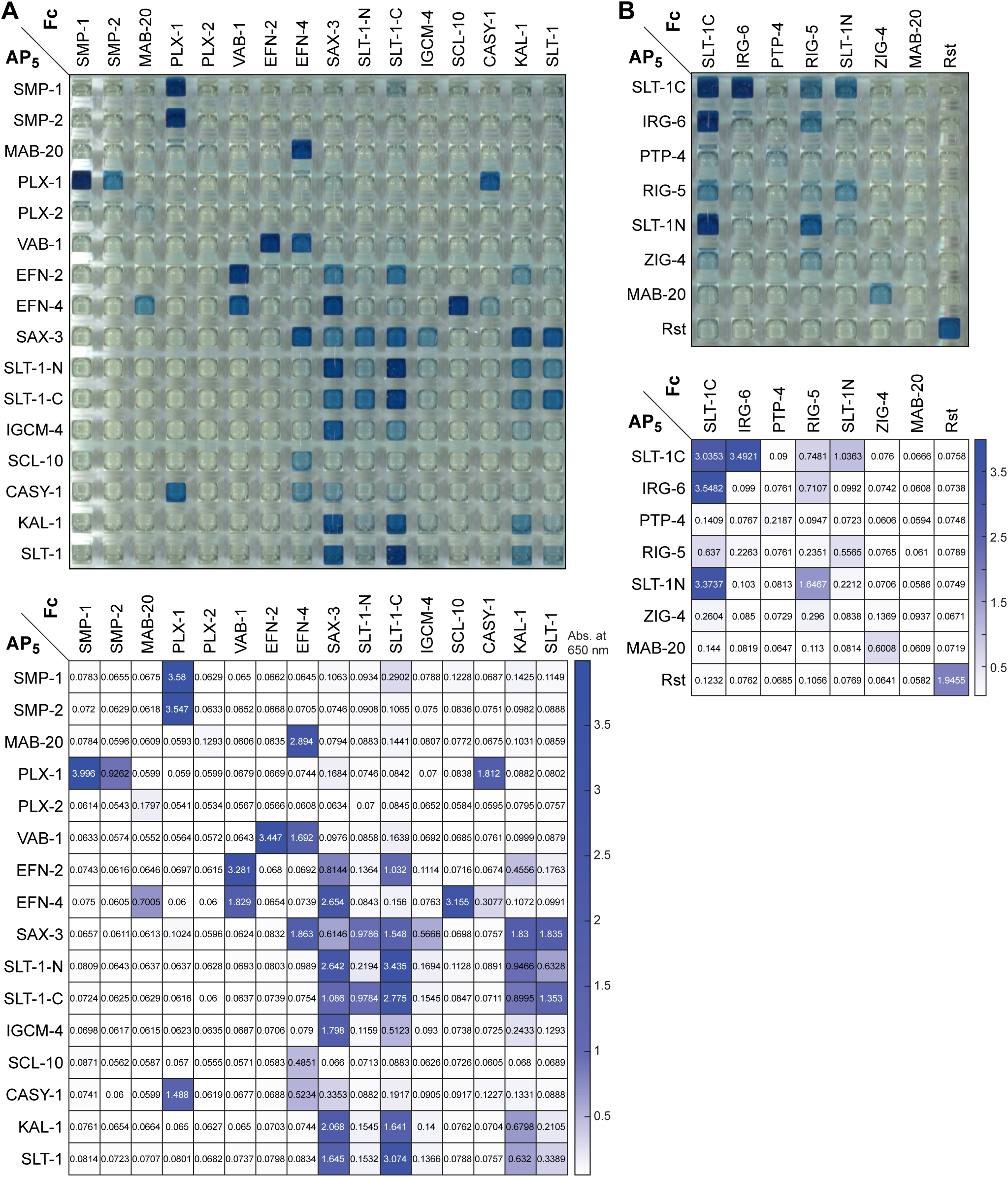
Interactions of axon guidance receptors and cues. **A.** Image of the 384-well plate and absorbance at 650 nm for the ECIA experiment for selected axon guidance-related proteins in Figure 3B. **B.** ECIA experiment for other guidance-related proteins. *D. melanogaster* Rst is a homodimeric protein and serves as a control.

**Figure S4.**
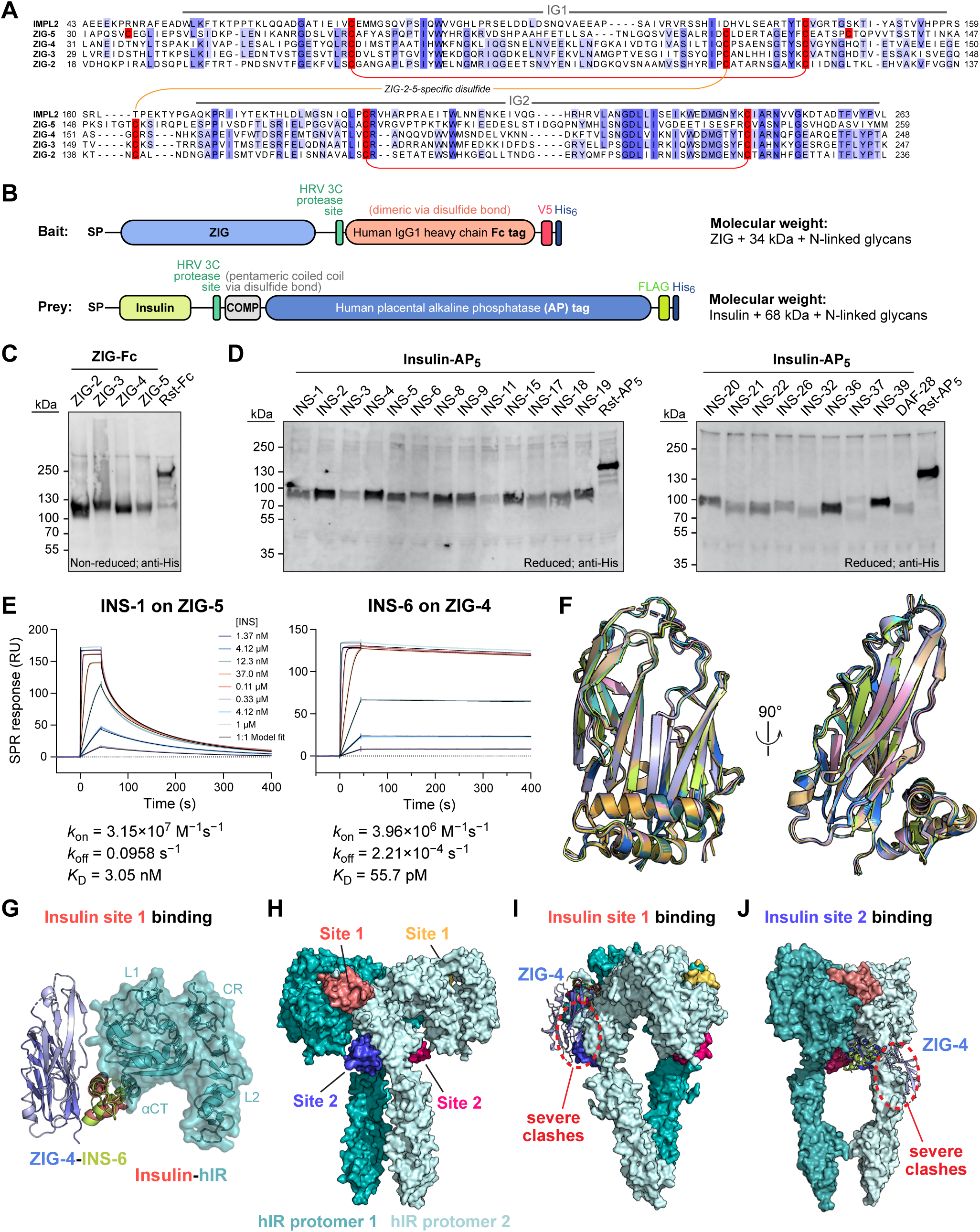
The ZIG-insulin interactome. **A.** Sequence alignment of four ZIGs and the fly ortholog, ImpL2. ZIG-2 to -5 carry a disulfide unique to all worm ZIGs. **B.** The ECIA construct design where ZIGs are depicted as bait and insulins as prey, as used in the experiment presented in Figure 4B. **C, D.** Expression of all insulin and ZIG constructs used in the experiment presented in Figure 4B. Expression of bait is shown in C and expression of prey in D. E. Kinetic fitting of SPR sensorgrams from Figure 4D with parameters. F. Superposition of three ZIG-4–INS-6 structures solved using three different crystal forms. G. The INS-6–ZIG-4 complex is compatible with insulins interacting with the L1 domains + αCT helix in insulin receptors. hIR: human insulin receptor; PDB ID: 3W11. H. Structure of the active T-like IR_2_-insulin_4_ structure from PDB ID: 6PXV. Four insulin-binding sites are shown in red, yellow, blue and pink. **I, J.** Insulin-bound ZIG-4 would severely clash with dimeric IR, regardless of insulin binding to site 1 (I), or site 2 (J).

**Figure S5.**
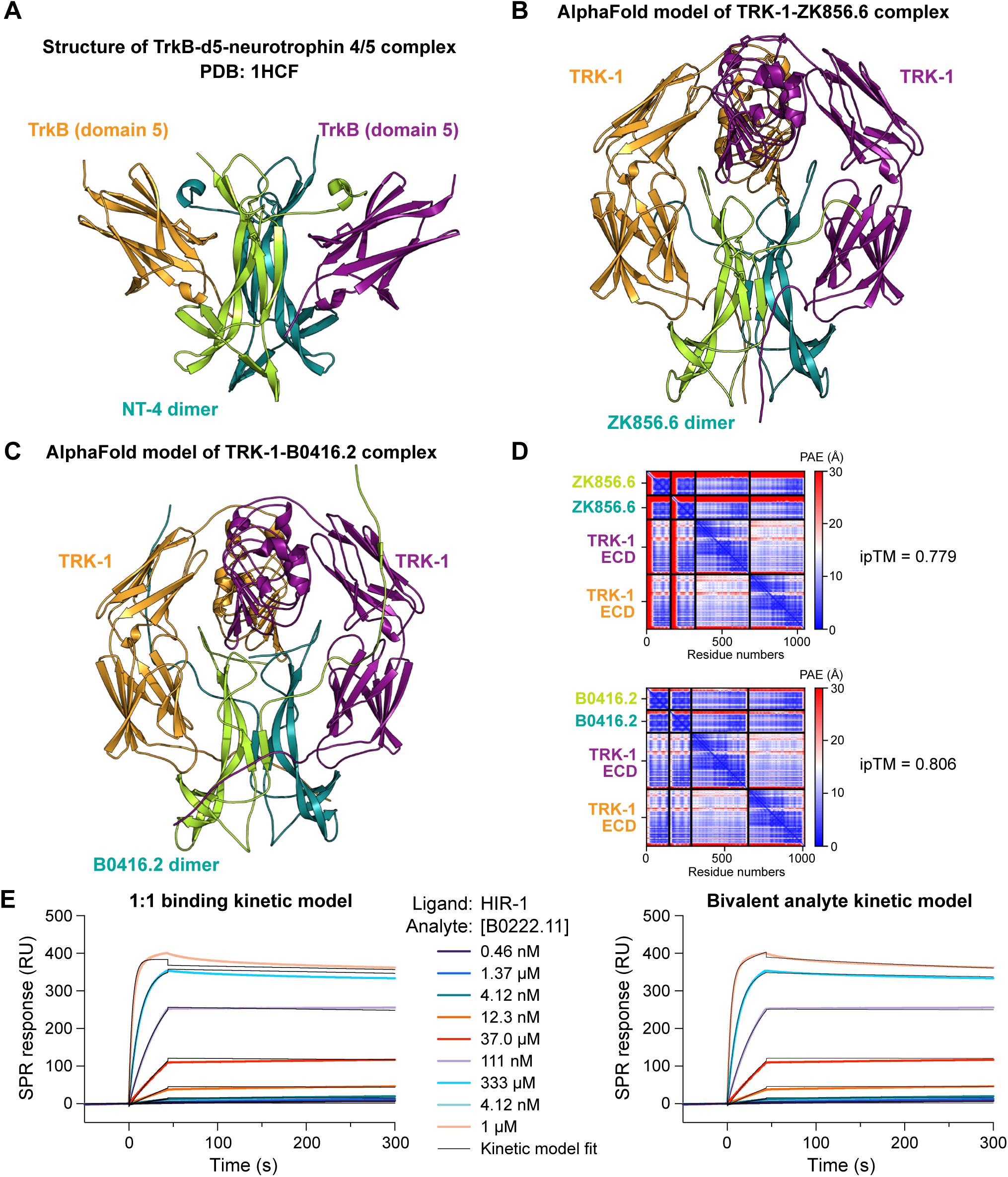
Comparison of AlphaFold models of complexes discovered by the ECIA screen with the structure of human ligand-bound neurotrophin receptor. A. Structure of human neurotrophin receptor, TrkB (domain 5) bound to NT4/5 (PDB: 1HCF). B. AlphaFold-predicted TRK-1 ectodomain bound to ZK856.6 at a 2:2 stoichiometry. C. AlphaFold-predicted TRK-1 ectodomain bound to B0416.2 at a 2:2 stoichiometry. D. PAE (Predicted Aligned Error) plots corresponding to models shown in B. and C. High ipTM (interface predicted Template Modelling) scores indicate high-confidence predictions. E. Kinetic fitting of SPR sensorgrams collected for the binding of B0222.11 to HIR-1, shown in Figure 5C.

**Figure S6.**
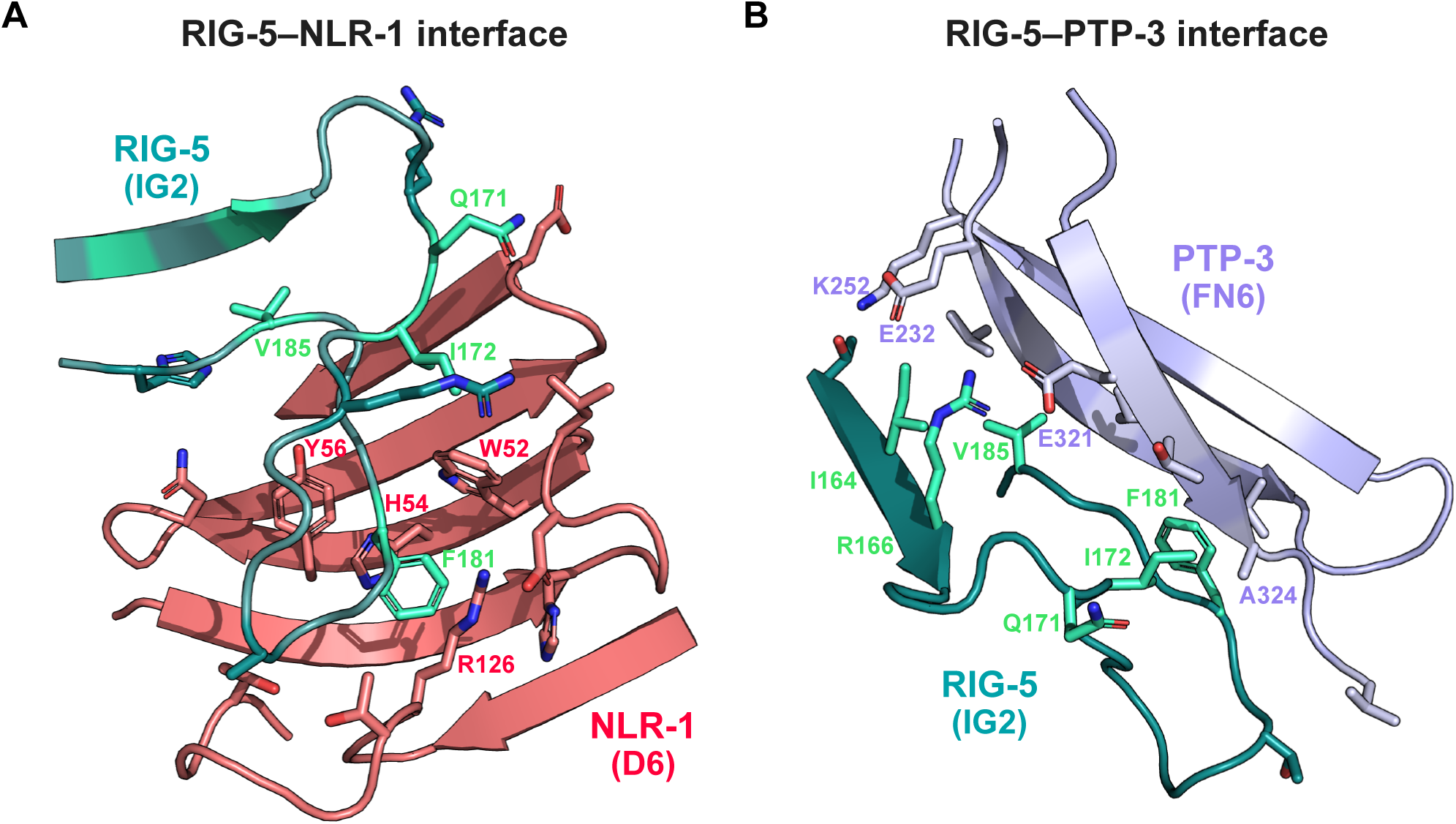
Interfaces observed in AlphaFold models of RIG-5-NLR-1 and RIG-5-PTP-3 complexes. **A.** The AlphaFold-predicted interface of RIG-5 (ECD) bound to NLR-1 (D6). **B.** The AlphaFold-predicted interface of RIG-5 (ECD) bound to PTP-3 (FN4-6). The RIG-5 residues mutated in the experiment presented in Figure 6H are shown in light cyan in A and B.

**Figure S7.**
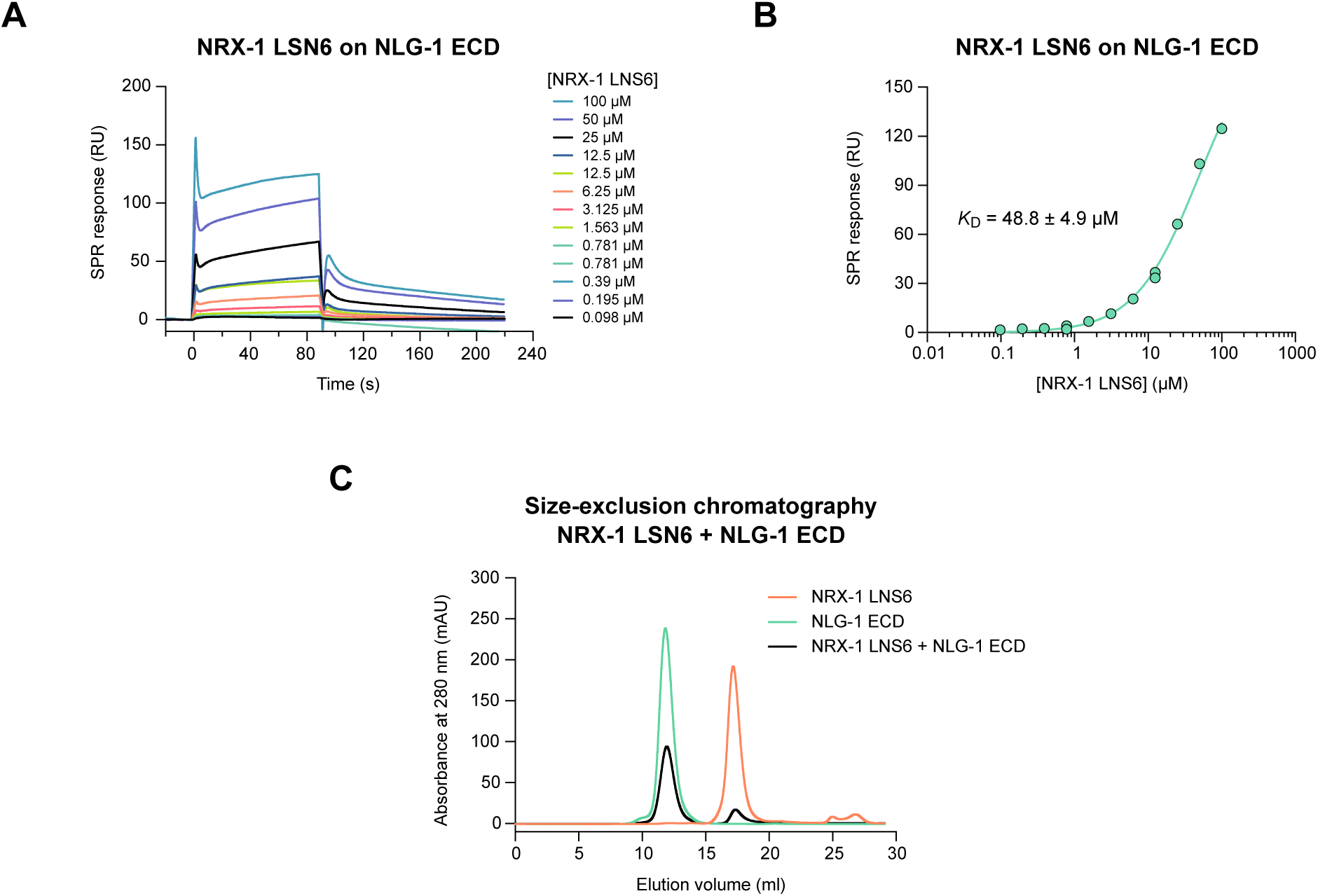
Binding experiments for NLG-1–NRX-1 complex. A. SPR sensorgrams for soluble NRX-1 LNS6 domain binding to immobilized NLG-1 ECD. B. Binding isotherm and *K*_D_ for binding shown in A. C. Size-exclusion chromatography runs for NRX-1 LNS-6 (orange), NLG-1 ECD (green) and the mixed sample (black).

**Table S1. Excel file containing even more data too large to fit in a PDF.** Ectodomains used in the interactome study by gene, transcript and protein names, sequence, domain composition, signal peptide and membrane anchoring predictions. TM: transmembrane. Relative expression levels are measured and reported in columns P and Q for bait and prey constructs, respectively.

**Table S2. Excel file containing even more data too large to fit in a PDF.**

**A.** Symmetrized *z*-scores using the MaxEnt method.

**B.** Asymmetric *z*-scores using the MaxEnt method.

**Table S3. Excel file containing even more data too large to fit in a PDF.**

List of interactions observed in the high-throughout ECIA experiment using our new MaxEnt method with 2-hour absorbance measurements. Interactions with only one orientation with *z*>3 are labeled pink in column G. For comparison, scoring according to our old method, geometric mean of trimmed z-scores (√(*z*_1_×*z*_2_)_old_) (Özkan, et al. *Cell*, 2013), are given in H, where a score of >20 was considered significant. Column I reports if the interaction or an orthologous one was reported before, based on a literature search. Alphafold-multimer (Colabfold version 1.5.2) iPTM scores for a subset of interacting pairs are in column J.

**Table S4. Excel file containing even more data too large to fit in a PDF.**

**A.** Canonical neighbors for every ectodomain tested, protein/sequence names in top row in bold.

**B.** All neighbors for every ectodomain tested, protein/sequence names in top row in bold.

**Table S5.** Data and refinement statistics for x-ray crystallography of the ZIG-4–INS-6 complex.

|  | **Tetragonal form** | **C-centered monoclinic** | **Primitive monoclinic** |
| --- | --- | --- | --- |
| **Data Collection** |  |  |  |
| Beamline | APS 24-ID-E | APS 24-ID-E | APS 24-ID-E |
| Space Group | $P4_22_12$ | $C2$ | $P2_1$ |
| *Cell Dimensions* |  |  |  |
| $a, b, c$ (Å) | 74.528, 74.528, 107.058 | 166.430, 56.654, 73.685 | 73.839, 55.739, 149.945 |
| $\alpha, \beta, \gamma$ (°) | 90, 90, 90 | 90, 113.247, 90 | 90, 93.775, 90 |
| Resolution (Å)* | 200-1.30 (1.38-1.30) | 200-2.30 (2.44-2.30) | 200-2.35 (2.49-2.35) |
| $R_{\text{sym}}$ (%) | 4.6 (120.1) | 7.1 (113.1) | 6.2 (124.8) |
| $\langle I \rangle / \langle \sigma(I) \rangle$ | 24.0 (1.78) | 9.7 (1.1) | 10.7 (0.9) |
| $CC_{1/2}$ | 0.999 (0.742) | 0.998 (0.652) | 0.998 (0.463) |
| Completeness (%) | 99.8 (99.4) | 98.4 (96.7) | 97.9 (97.1) |
| Redundancy | 12.8 (12.2) | 3.4 (3.3) | 3.1 (3.1) |
| **Refinement** |  |  |  |
| Resolution (Å)* | 50-1.30 (1.32-1.30) | 53.13-2.30 (2.38-2.30) | 74.81-2.35 (2.60-2.50) |
| Reflections | 73,784 | 27,969 | 50,096 |
| $R_{\text{cryst}}$ (%) | 14.47 (27.19) | 21.52 (43.52) | 19.66 (38.72) |
| $R_{\text{free}}$ (%)** | 16.89 (31.58) | 25.18 (47.21) | 23.98 (44.33) |
| *Number of atoms* |  |  |  |
| Protein | 2,128 | 3,878 | 7,755 |
| Ligand/Glycans | 11 | 0 | 0 |
| Water | 305 | 7 | 39 |
| *Average B-factors (Å<sup>2</sup>)* |  |  |  |
| All | 24.8 | 76.2 | 83.1 |
| Protein | 23.0 | 76.3 | 83.2 |
| Ligand | 36.3 | N/A | N/A |
| Solvent | 37.0 | 60.2 | 57.5 |
| *R.m.s. deviations from ideality* |  |  |  |
| Bond Lengths (Å) | 0.012 | 0.003 | 0.008 |
| Bond Angles (°) | 1.291 | 0.631 | 1.002 |
| *Ramachandran plot* |  |  |  |
| Favored (%) | 98.41 | 96.67 | 97.24 |
| Outliers (%) | 0.40 | 0.00 | 0.00 |
| Rotamer Outliers (%) | 0.43 | 0.48 | 0.39 |
| All-atom Clashscore<sup>‡</sup> | 3.34 | 3.31 | 3.91 |

\* The values in parentheses are for reflections in the highest resolution bin.
\*\* 5% of reflections (3,747) for tetragonal crystals, 5% of reflections (1,373) for C-centered monoclinic crystals, and 4% of reflections (1,977) for primitive monoclinic crystals were not used during refinement for cross validation purposes.
<sup>‡</sup> Clashscores were calculated by *phenix.refine* (Phenix version 1.21).
N/A: Not applicable.

**Table S6. Excel file containing even more data too large to fit in a PDF.** Experimental details and parameters for all surface plasmon experiments included in the manuscript. Biacore chips are purchased from Cytiva. HBSp+: 10 mM HEPES, 150 mM NaCl, 0.05% Tween-20.

**Table S7. Excel file containing even more data too large to fit in a PDF.**

185 experimental PPIs based on the number of chemical synapses associated with each interaction. Interactions where there was no expression data for one or both of the binding partners are labeled N/A. We randomize the neuron connectome as a random control.

